# Polyunsaturated fatty acids inhibit a pentameric ligand-gated ion channel through one of two binding sites

**DOI:** 10.1101/2021.10.08.463634

**Authors:** Noah M. Dietzen, Mark J. Arcario, Lawrence J. Chen, John T. Petroff, Kathiresan Krishnan, Grace Brannigan, Douglas F. Covey, Wayland W. L. Cheng

## Abstract

Polyunsaturated fatty acids (PUFAs) inhibit pentameric ligand-gated ion channels (pLGICs) but the mechanism of inhibition is not well understood. The PUFA, docosahexaenoic acid (DHA), inhibits agonist responses of the pLGIC, ELIC, more effectively than palmitic acid, similar to the effects observed in the GABA_A_ receptor and nicotinic acetylcholine receptor. Using photo-affinity labeling and coarse-grained molecular dynamics simulations, we identified two fatty acid binding sites in the outer transmembrane domain (TMD) of ELIC. Fatty acid binding to the photolabeled sites is selective for DHA over palmitic acid, and specific for an agonist-bound state. Hexadecyl-methanethiosulfonate modification of one of the two fatty acid binding sites in the outer TMD recapitulates the inhibitory effect of PUFAs in ELIC. The results demonstrate that DHA selectively binds to multiple sites in the outer TMD of ELIC, but that state-dependent binding to a single intrasubunit site mediates DHA inhibition of ELIC.

## INTRODUCTION

Fatty acids are major components of the cell membrane, and modulators of many ion channels including pentameric ligand-gated ion channels (pLGICs) (1–6). Fatty acids inhibit neuronal pLGICs including the GABA_A_ receptor (GABA_A_R) (4) and nicotinic acetylcholine receptor (nAchR) (5,6), by reducing peak channel responses to agonist. Polyunsaturated fatty acids (PUFAs) have a stronger inhibitory effect than mono- or saturated fatty acids (5,6). While the physiologic significance of PUFA inhibition of pLGICs is not fully understood, neuronal PUFAs affect neurologic processes in which pLGIC function is critical such as neurodevelopment (7) and cognition (8). Neuronal PUFA content can change dramatically in pathologic conditions such as stroke and seizure (9), and PUFA inhibition of pLGICs under these conditions is likely an important determinant of neuronal excitability (1,10).

Fatty acids modulate ion channel function allosterically by one of two general mechanisms: by direct binding to specific sites in the protein, or by altering the physical properties of the lipid bilayer and thereby indirectly impacting protein structure. There is evidence to support both mechanisms in different ion channels including pLGICs (11,12). A direct mechanism is thought to be important in pLGICs based on a crystal structure of *Gloeobacter* ligand-gated ion channel (GLIC) in complex with docosahexaenoic acid (DHA), which showed a single binding site for DHA in the outer portion of the transmembrane domain (TMD) of this channel (3). However, whether fatty acids bind to other sites in pLGICs, and whether these sites are specific for PUFAs and mediate PUFA inhibition of channel activity are not established. It is possible that fatty acids bind to other sites in GLIC that were not resolved in the crystal structure due to greater flexibility of the fatty acid within those sites. Mutagenesis studies and molecular dynamics simulations also support a direct mechanism of fatty acid modulation of large conductance calcium-activated potassium channels (BK) (13) and voltage-gated potassium channels (Kv) (14,15), but biochemical identification of fatty acid binding sites in these and other ion channels remains a challenge.

Photo-affinity labeling is an alternative approach to identify lipid and small molecule binding sites in membrane proteins. Fatty acid photolabeling reagents have been used previously to identify fatty acid binding proteins (16–18), but not to map binding sites at the residue level. We introduce a new photolabeling reagent with optimal photochemistry coupled with intact protein and middle-down mass spectrometry (MS) (19–21) to identify fatty acid binding sites in the pLGIC, *Erwinia* ligand-gated ion channel (ELIC) (22). Combining this approach with coarse-grained molecular dynamics (CGMD) simulations, we determine two fatty acid binding sites in the outer TMD of ELIC that are specific for DHA over palmitic acid when agonist is bound to the channel. Occupancy of one of these sites mediates the inhibitory effect of DHA on ELIC channel responses. The results argue for a direct mechanism of DHA inhibition of ELIC through state-dependent binding to a single site.

## RESULTS

### DHA inhibits ELIC channel function

ELIC has been a useful model channel for investigating the structural determinants of lipid (23,24) and anesthetic (25–27) modulation of pLGICs. We sought to determine if ELIC is a suitable model for examining the mechanism of fatty acid inhibition of pLGICs. ELIC was reconstituted in giant liposomes composed of a 2:1:1 ratio of POPC:POPE:POPG for excised patch voltage-clamp recordings and channel responses were evaluated by rapid application of the agonist, cysteamine, at 30 mM for 1 min. ELIC responses showed fast activation followed by a slower current decay or desensitization, consistent with previous reports of ELIC function in liposomes or HEK cells (24,28). The effect of fatty acids such as DHA (22:6) was determined by pre-applying fatty acid to the patch for 3 min followed by rapid application of cysteamine with the same concentration of fatty acid used in the pre-application (Figure 1A). Longer pre-application times with fatty acid did not yield a greater effect, indicating that 3 min was sufficient time to equilibrate the patch membrane with DHA from solution.

**Figure 1:**
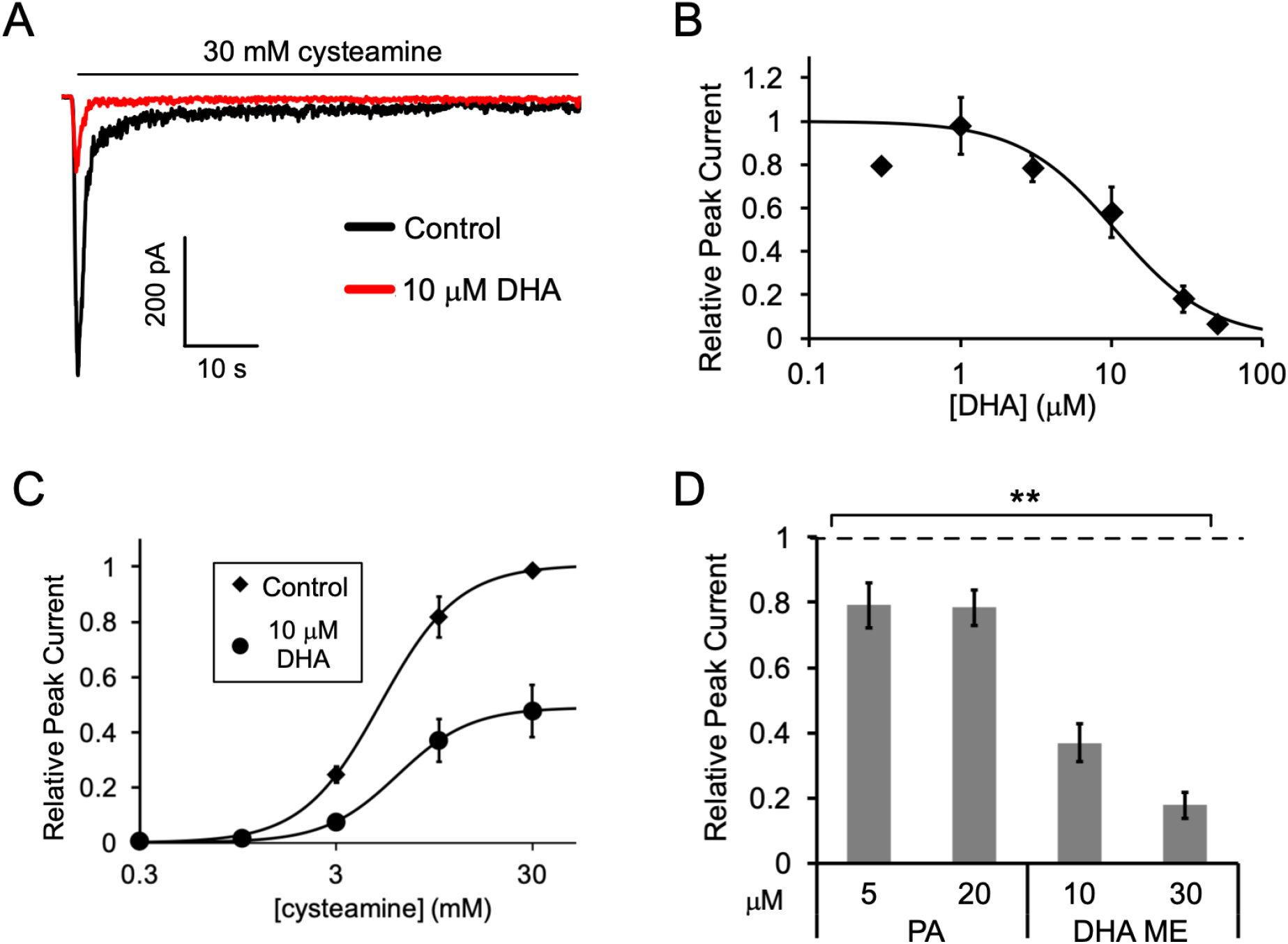
Fatty acid inhibition of ELIC. (A) Sample currents from excised patch-clamp (−60 mV) of WT ELIC in 2:1:1 POPC:POPE:POPG giant liposomes. Current responses are to 30 mM cysteamine before and after 3 min pre-application 10 μM DHA. (B) Peak current responses to 30 mM cysteamine normalized to control (absence of DHA) as a function of DHA concentration (n=4-8, 0.3 and 50 μM n=1, ±SEM). The data are fit to a sigmoidal function yielding an IC_50_ of 10.6 μM. (C) Peak current responses normalized to maximum response as a function of cysteamine concentration in the absence or presence of 10 μM DHA (n=7, ±SEM, control EC_50_ = 5.5±0.7 mM, 10 μM DHA EC_50_ = 6.2±0.6 mM). (D) Peak current responses to 30 mM cysteamine with 3 min pre-application of PA (palmitic acid) and DHA ME (DHA methyl ester) normalized to control (no lipid) (n=4-6, ±SEM, ** p<0.01 compared to control).

Pre-application of DHA caused ELIC peak responses to decrease with an IC_50_ of 10.6 μM (Figure 1A and 1B); there was almost complete inhibition of ELIC responses at 50 μM DHA. DHA also caused a ~50% reduction in steady state current relative to the peak current in the absence of DHA (IC_50_ = 9.8 μM) and a similar percent reduction in the rate of current decay (IC_50_ = 10.5 μM) (Figure 1—figure supplement 1A and 1B). However, the EC_50_ of cysteamine activation showed no significant change with the pre-application of 10 μM DHA (Figure 1C). Furthermore, pre-application with the saturated fatty acid, palmitic acid (PA, 16:0), significantly reduced ELIC peak responses to 30 mM cysteamine, but to a much lesser degree than DHA (Figure 1D). 5 and 20 μM PA yielded the same effect indicating that saturating concentrations of PA were used. Interestingly, the methyl ester of DHA (DHA ME) strongly inhibited ELIC similar to DHA (Figure 1D), indicating that the carboxylate DHA headgroup is not required for this inhibitory effect. The effects of fatty acids on ELIC are qualitatively similar to the effects reported in the GABA_A_R (4,5,12) and nAchR (6,29). PUFAs inhibit gating efficacy without shifting agonist potency, and DHA ME is just as effective as DHA (10). The results suggest that the mechanism of fatty acid inhibition of the GABA_A_R and nAchR is also present in ELIC.

**Figure 1—figure supplement 1:**
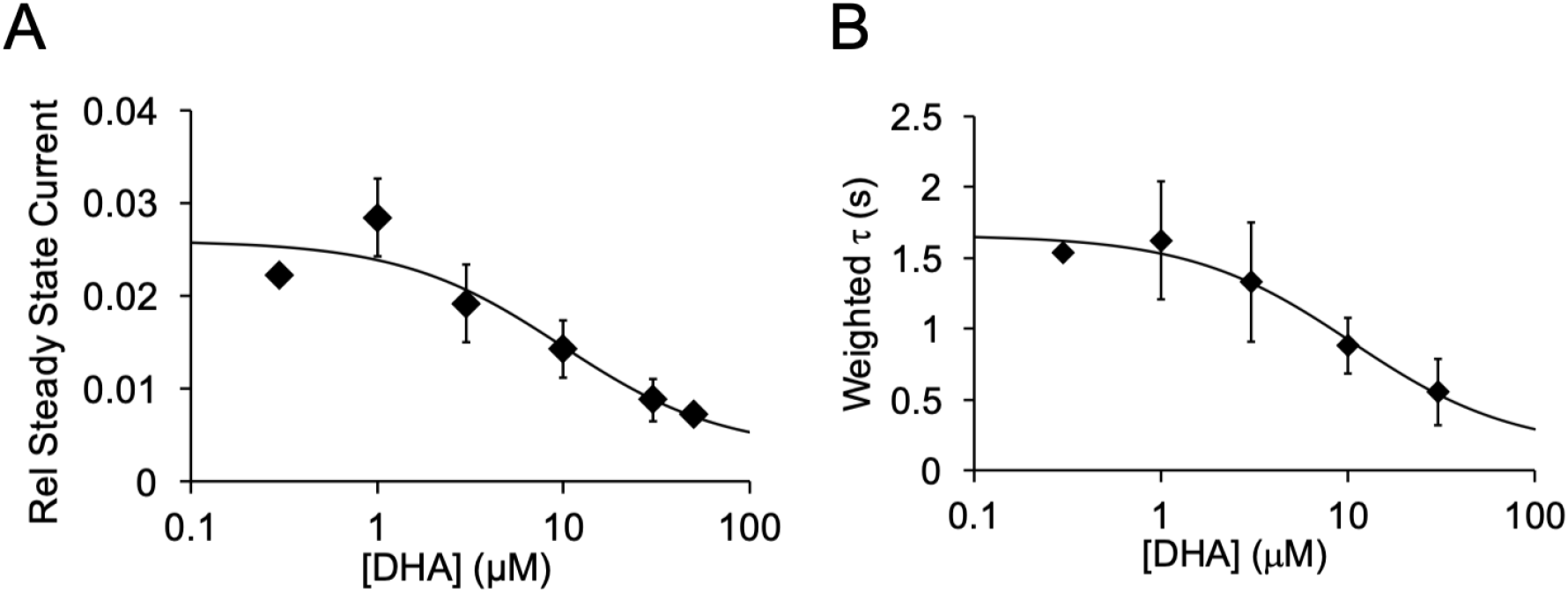
(A) Steady state current (60 s after exposure to 30 mM cysteamine) with varying concentrations of DHA normalized to peak current in the absence of DHA. The data are fit to a sigmoidal function yielding an IC_50_ of 9.8 μM (n=4-8, 0.3 and 50 μM n=1, ±SEM). (B) Weighted time constant (τ) of current decay for responses to 30 mM cysteamine at varying concentrations of DHA (IC_50_ = 10.5 μM) (n=4-8, 0.3 and 50 μM n=1, ±SEM).

Two possible explanations for why DHA is a stronger inhibitor of ELIC than PA are that the two fatty acids bind to different sites in ELIC or that PA has a lower affinity for the same binding site as DHA. To begin examining the interaction of fatty acids with ELIC, we performed CGMD on a single ELIC pentamer in a 2:1:1 POPC:POPE:POPG membrane containing either 4 mol% PA or DHA. CGMD allows for a significant amount of lipid diffusion over the simulation time scale, which permits increased sampling of protein-lipid interactions. We first quantified lipid sorting using the boundary lipid metric, *B* (Equation 2), where *B* > 1 indicates enrichment compared to bulk membrane and *B* < 1 indicates relative depletion (30). DHA shows significant enrichment in the lipids surrounding ELIC (B = 2.43 ± 0.16, ±SEM, n=4), while PA shows a relative depletion (B = 0.51 ± 0.02, ±SEM, n=4). To identify localized density that would indicate specific binding, we calculated the two-dimensional radial enhancement for each fatty acid species with ELIC centered at the origin (Figure 2). In these plots, white tiles represent no enrichment over expected bulk membrane density (radial enhancement, 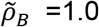); increasing enrichment is represented by deepening red tiles 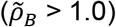 and increasing depletion is represented by deepening blue tiles 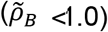. It is immediately apparent that DHA shows several localized areas of high enrichment indicating specific sites of binding to ELIC, whereas PA shows areas of minimal enrichment that are more diffuse suggesting no specific binding (Figure 2). Binding of DHA is observed mainly in two intrasubunit grooves located between M4 and M1 or M3. Moreover, these areas of localized binding are noted mostly in the outer leaflet (Figure 2, top right), suggesting DHA acts through a specific binding site in the outer leaflet of the membrane. These results are consistently demonstrated over multiple simulation replicates (Figure 2—figure supplement 1; Figure 2—figure supplement 2), indicating that that in 2:1:1 POPC:POPE:POPG lipid membranes in which ELIC responses were measured, DHA binds to specific intrasubunit sites in the outer leaflet, which PA is not seen to strongly interact with. This supports the notion that DHA inhibits ELIC by binding to a specific site, while PA has low affinity for ELIC.

**Figure 2:**
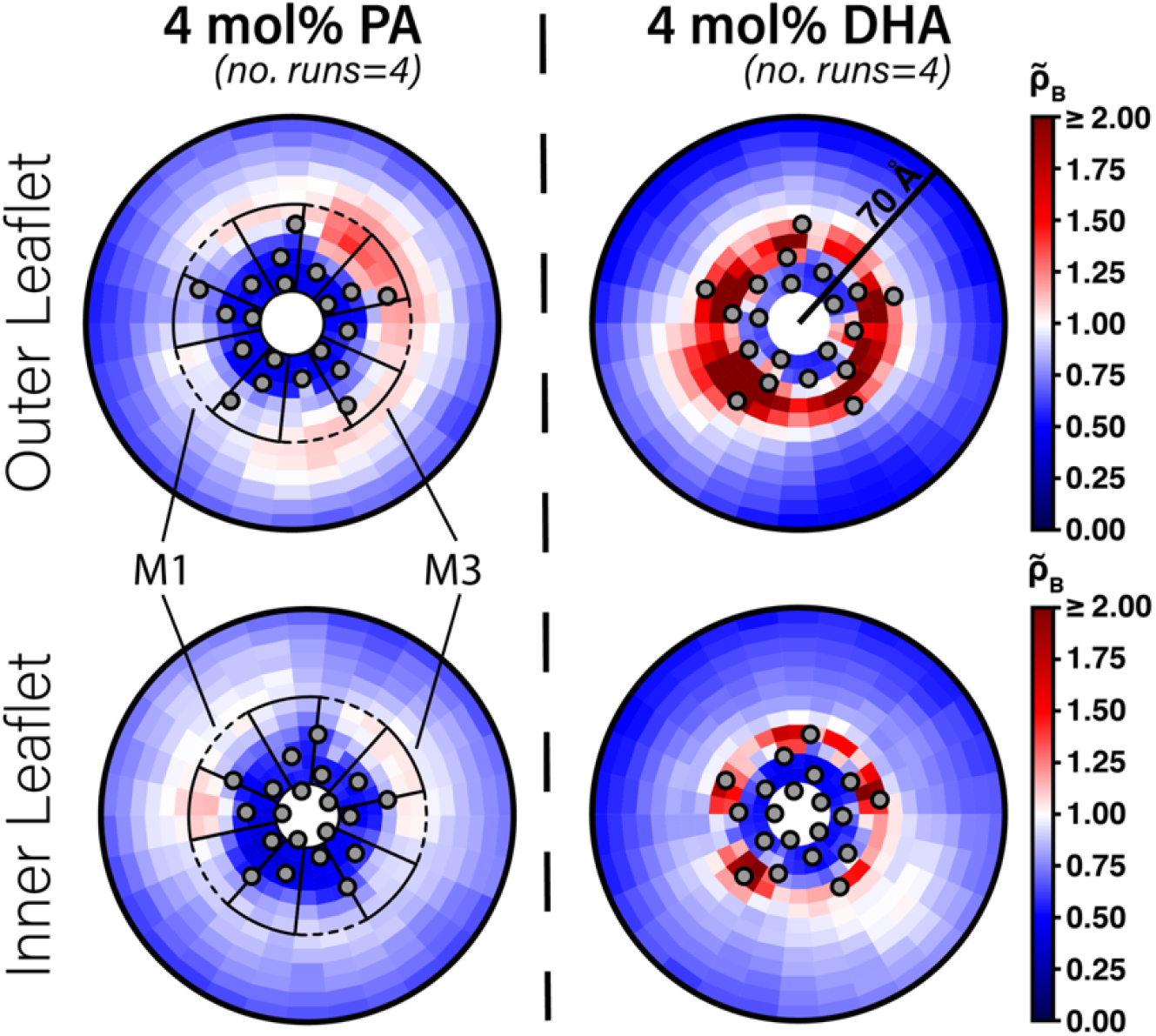
Enrichment of PA and DHA in ELIC. Two-dimensional enrichment plots for the 4 mol% PA (left column) and 4 mol% DHA (right column) simulation conditions are shown for fatty acid species within 70 Å of the ELIC pore. Separate enrichment for the outer (top row) and inner (bottom row) leaflet are plotted with the color bar at right demonstrating relative enrichment values compared to bulk membrane. 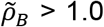 indicates enrichment over bulk membrane and 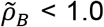 indicates relative depletion compared to bulk membrane (Equation 4). Data presented in this figure represent the average across four simulation runs. The gray circles in each plot represent the location of the transmembrane helices relative to the ELIC pore. The dashed black outline sectors demonstrate the boundaries of the M1 site as used for density affinity threshold calculations, whereas the solid black outline sectors demonstrate the boundaries of the M3 site.

**Figure 2—figure supplement 1:**
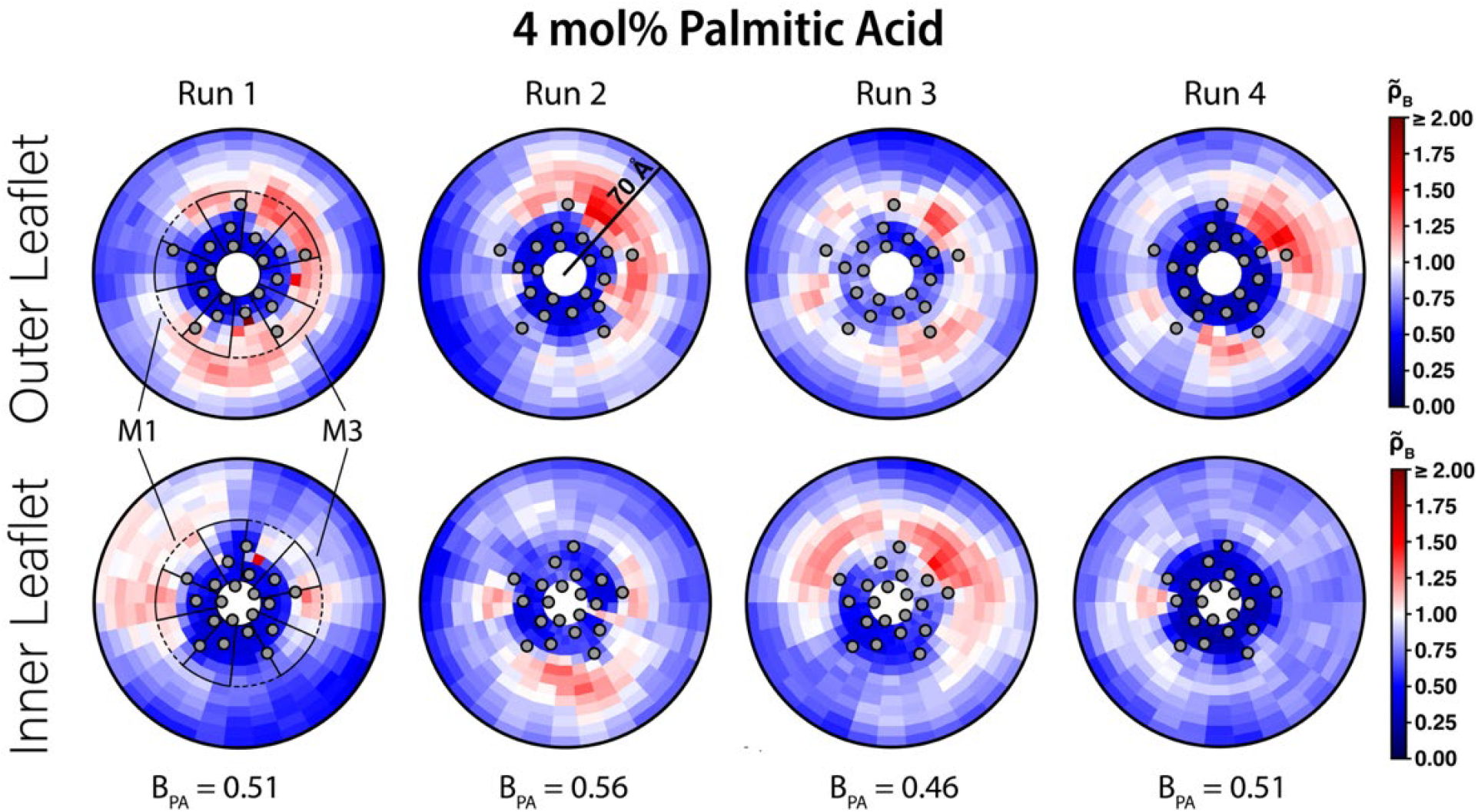
Two-dimensional enrichment plots for the four individual 4 mol% PA simulations showing enrichment in the outer (top row) and inner (bottom row) leaflets within 70 Å of the ELIC pore. The boundary lipid metric, *B* (Equation 2), is given for each individual CGMD simulation run underneath the two-dimensional enrichment plot with the average (±SEM) across all four runs being 0.51 ± 0.04. The gray circles in each plot represent the location of the transmembrane helices relative to the ELIC pore. The dashed black outline sectors demonstrate the boundaries of the M1 site as used for density affinity threshold calculations, whereas the solid black outline sectors demonstrate the boundaries of the M3 site. The color bar for relative enrichment and depletion is placed at the right side of the figure.

**Figure 2—figure supplement 2:**
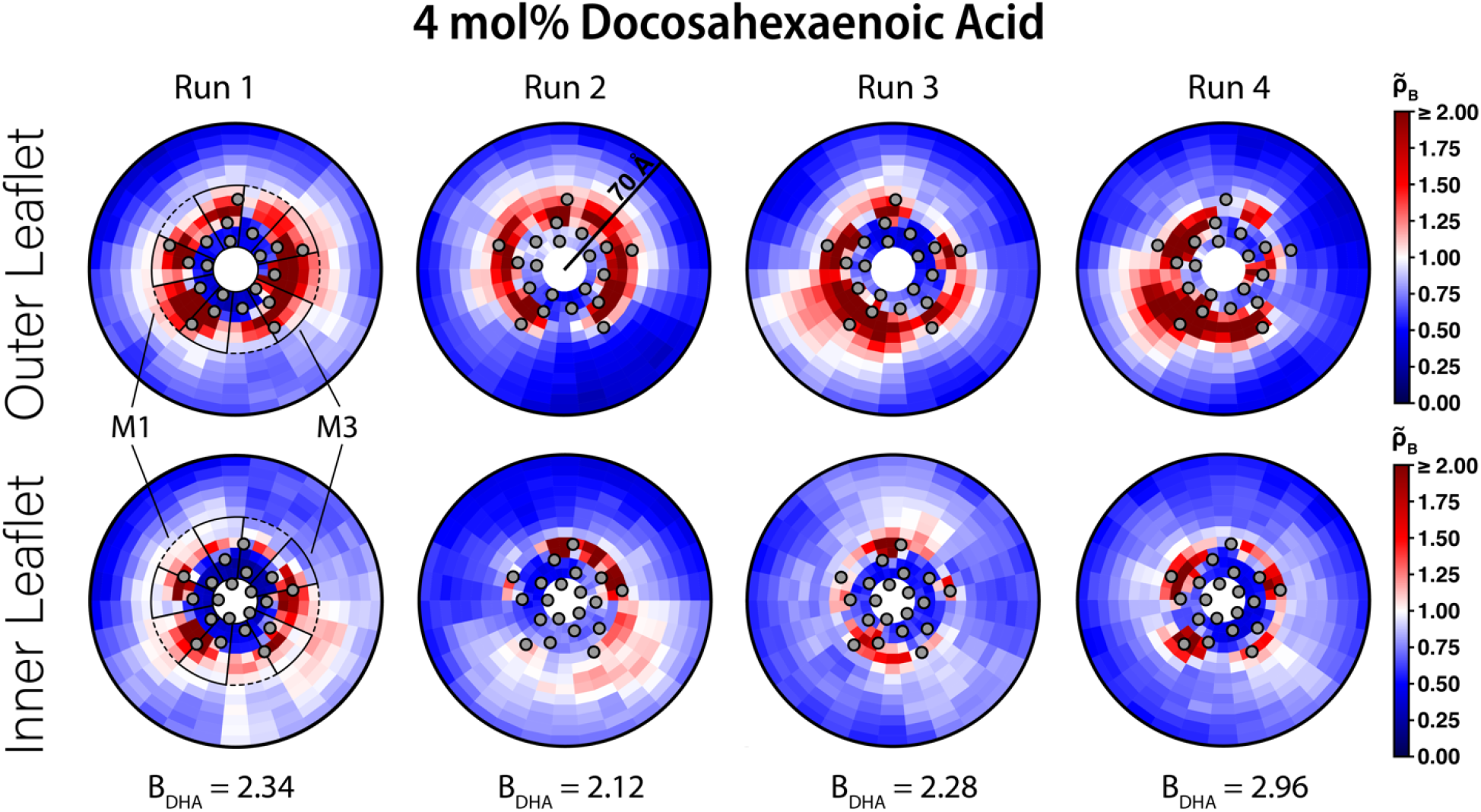
Two-dimensional enrichment plots for the four individual 4 mol% DHA simulations showing enrichment in the outer (top row) and inner (bottom row) leaflets within 70 Å of the ELIC pore. The boundary lipid metric, *B* (Equation 2), is given for each individual CGMD simulation run underneath the two-dimensional enrichment plot with the average (±SEM) across all four runs being 2.43 ± 0.16. The gray circles in each plot represent the location of the transmembrane helices relative to the ELIC pore. The dashed black outline sectors demonstrate the boundaries of the M1 site as used for density affinity threshold calculations, whereas the solid black outline sectors demonstrate the boundaries of the M3 site. The color bar for relative enrichment and depletion is at the right side of the figure.

### Characterization of a fatty acid photolabeling reagent, KK242, in ELIC

To further identify fatty acid binding sites in ELIC, we used photo-affinity labeling coupled with MS. We began by testing a commercially available fatty acid photolabeling reagent, 9-(3-pent-4-ynyl-3-H-diazirin-3-yl)-nonanoic acid or pacFA (Avanti Polar Lipids, Figure 3A). This bifunctional reagent contains a photo-reactive aliphatic diazirine in the alkyl tail and an alkyne for click chemistry (17). We photolabeled purified ELIC with 100 μM pacFA and analyzed the photolabeled protein by intact protein MS (20,21,31,32). Even at this high concentration, no photolabeling of ELIC by pacFA could be detected (Figure 3B).

**Figure 3:**
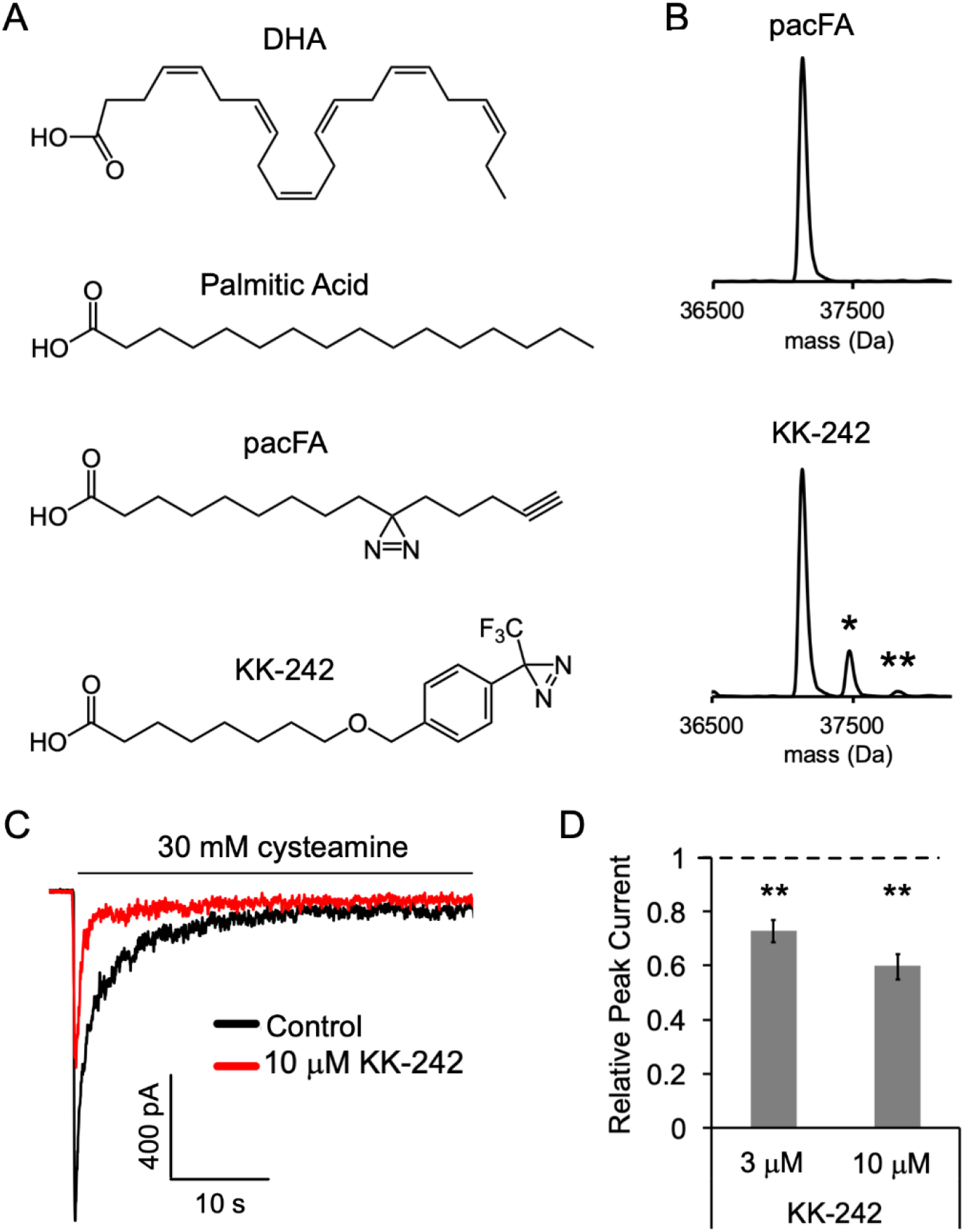
Fatty acid reagent photolabeling and inhibition of ELIC. (A) Chemical structures of DHA, PA, pacFA, and KK-242. (B) Deconvoluted intact protein mass spectra of WT ELIC photolabeled with 100 μM pacFA and 100 μM KK-242. In both spectra, the highest intensity peak corresponds to the WT ELIC subunit, and in the KK-242 spectrum, the peaks labeled * and ** indicate an ELIC subunit with 1 and 2 KK-242 adducts, respectively. (C) Sample currents from excised patch-clamp (−60 mV) of WT ELIC in 2:1:1 POPC:POPE:POPG giant liposomes. Current responses are to 30 mM cysteamine before and after 3 min pre-application 10 μM KK-242. (D) Peak current responses to 30 mM cysteamine normalized to control (absence of KK-242) as a function of KK-242 concentration (n=4, ±SEM).

We have previously shown that photolabeling reagents that contain an aliphatic diazirine almost exclusively label nucleophilic amino acid side chains such as glutamate or aspartate (21). Since the alkyl tail of pacFA, where the photo-reactive diazirine is located, is expected to interact with the hydrophobic regions of the ELIC TMD, one might expect that pacFA will not photolabel ELIC as there are no strong nucleophiles in the TMD facing the hydrophobic core of the membrane. In contrast to an aliphatic diazirine, a trifluoromethylphenyl-diazirine (TPD) shows less selectivity between amino acid side chains and offers favorable photochemistry for photolabeling studies with lipids (21,32). Thus, we synthesized a fatty acid analogue photolabeling reagent, KK-242, which contains an ether-linked TPD group in the fatty acid alkyl tail and approximates the molecular dimensions of PA (~20 Å in length) (Figure 3A). Intact protein MS of ELIC photolabeled with 100 μM KK-242 showed two additional peaks corresponding to the mass of an ELIC subunit with one and two KK-242 labels (labeling efficiency ~15%) (Figure 3B). This result indicates that KK-242 labels ELIC efficiently, and that there is a minimum of two non-overlapping labeled sites per subunit.

We next verified that this fatty acid analogue also inhibits ELIC function. 3 and 10 μM KK-242 inhibited ELIC peak responses (Figure 3C and 3D) with similar efficacy, indicating that these concentrations are near saturation. The efficacy of KK-242 was slightly higher than PA, but less than DHA. KK-242 also modestly decreased relative steady state current and the time constant of current decay (Figure 3—figure supplement 1). Thus, KK-242 is a suitable fatty acid analogue photolabeling reagent, reproducing the inhibitory effects of fatty acids on ELIC responses.

**Figure 3—figure supplement 1:**
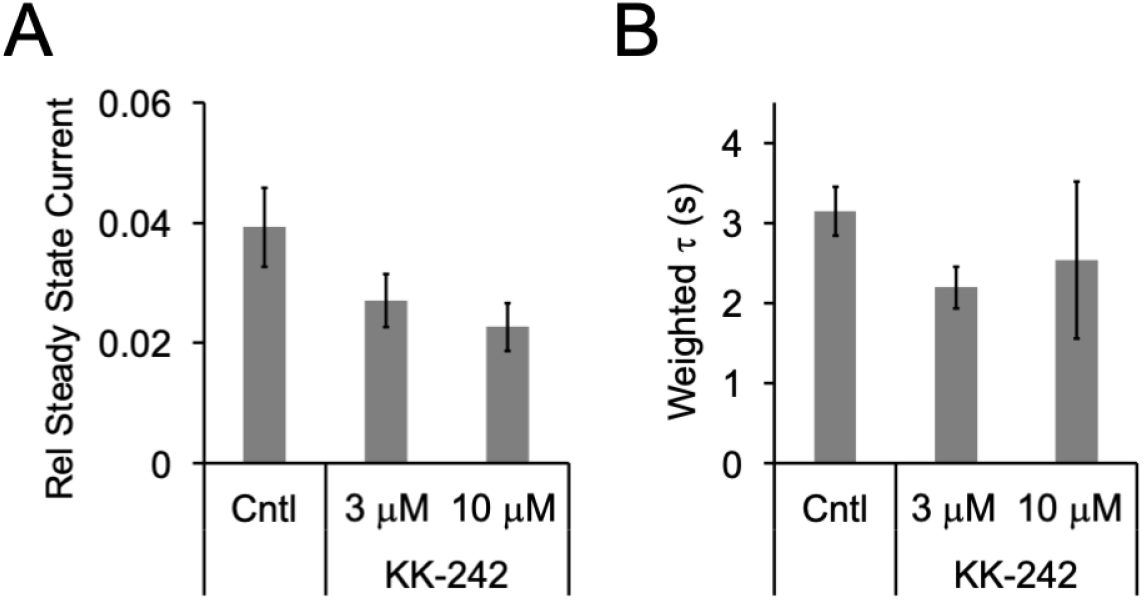
(A) Steady state current (60 s after exposure to 30 mM cysteamine) in the absence or presence of KK-242 (3 and 10 μM) normalized to peak current in the absence of KK-242 (n=4, ±SEM). (B) Weighted time constant (τ) of current decay for responses to 30 mM cysteamine in the absence or presence of KK-242 (3 and 10 μM) (n=4, ±SEM).

### DHA binds to two sites in ELIC in an agonist-bound state

To identify the photolabeled sites of KK-242 in ELIC, we analyzed ELIC photolabeled with 100 μM KK-242 with and without 30 mM cysteamine using a tryptic middle-down MS approach (21). Initial analysis of the LC-MS data using PEAKS showed 100% coverage of the ELIC sequence, including high intensity peptides for all four transmembrane helices and KK-242 modified peptides encompassing M3 and M4 (Figure 4—figure supplement 1). Both photolabeled peptides had a longer retention time than the corresponding unlabeled peptide, as would be expected with reverse phase chromatography (Figure 4—figure supplement 2), and high mass accuracy (Figure 4—table supplement 1). No photolabeled peptides were identified in the extracellular domain, M1 or M2.

The labeling efficiencies of ELIC subunits at 100 μM KK-242 with and without 30 mM cysteamine were indistinguishable by intact protein MS (~15%). Labeling efficiencies of M4 and M3 with and without cysteamine, and the corresponding fragmentation spectra were also similar (Figure 4—figure supplement 3). Thus, we will focus on the fragmentation spectra of photolabeled ELIC in the presence of cysteamine. Manual analysis of the MS2 spectra of both photolabeled peptides revealed fragment ions containing the KK-242 mass (Figure 4A and 4B). For the M3 peptide, three *b* ions localize the photolabeled residue to Q264 (Figure 4A). For the M4 peptide, a series of *b* and *y* ions indicate two photolabeled residues: C313 and R318 (Figure 4B). In both peptides, the same fragment ions with and without the KK-242 mass are frequently present most likely representing partial neutral loss of KK-242. While the possibility of 100% of neutral loss raises some uncertainty about the residues that are photolabeled, the spectra are most consistent with photolabeling of Q264 in M3, and C313 and R318 in M4. Q264 is located in M3 on one side of M4, while C313 is facing the other side of M4 adjacent to M1 (Figure 4A and 4B). Most structures of ELIC do not resolve R318 at the C-terminal end of M4; however, a recent cryo-EM structure of ELIC shows this residue facing M3 adjacent to Q264 (33). We docked KK-242 in two volumes that encompass the photolabeled residues and the outer interfacial region of the ELIC TMD. While many binding poses were obtained (KK-242 has 13 rotatable bonds), Figure 4A and 4B show the top pose from each site where the diazirine is closest to the photolabeled residue. When this is the case, the carboxylate group of KK-242 is approximately within the interfacial region of the membrane (Figure 4A and 4B). Thus, KK-242 photolabels two distinct sites on either side of outer portion of M4.

**Figure 4:**
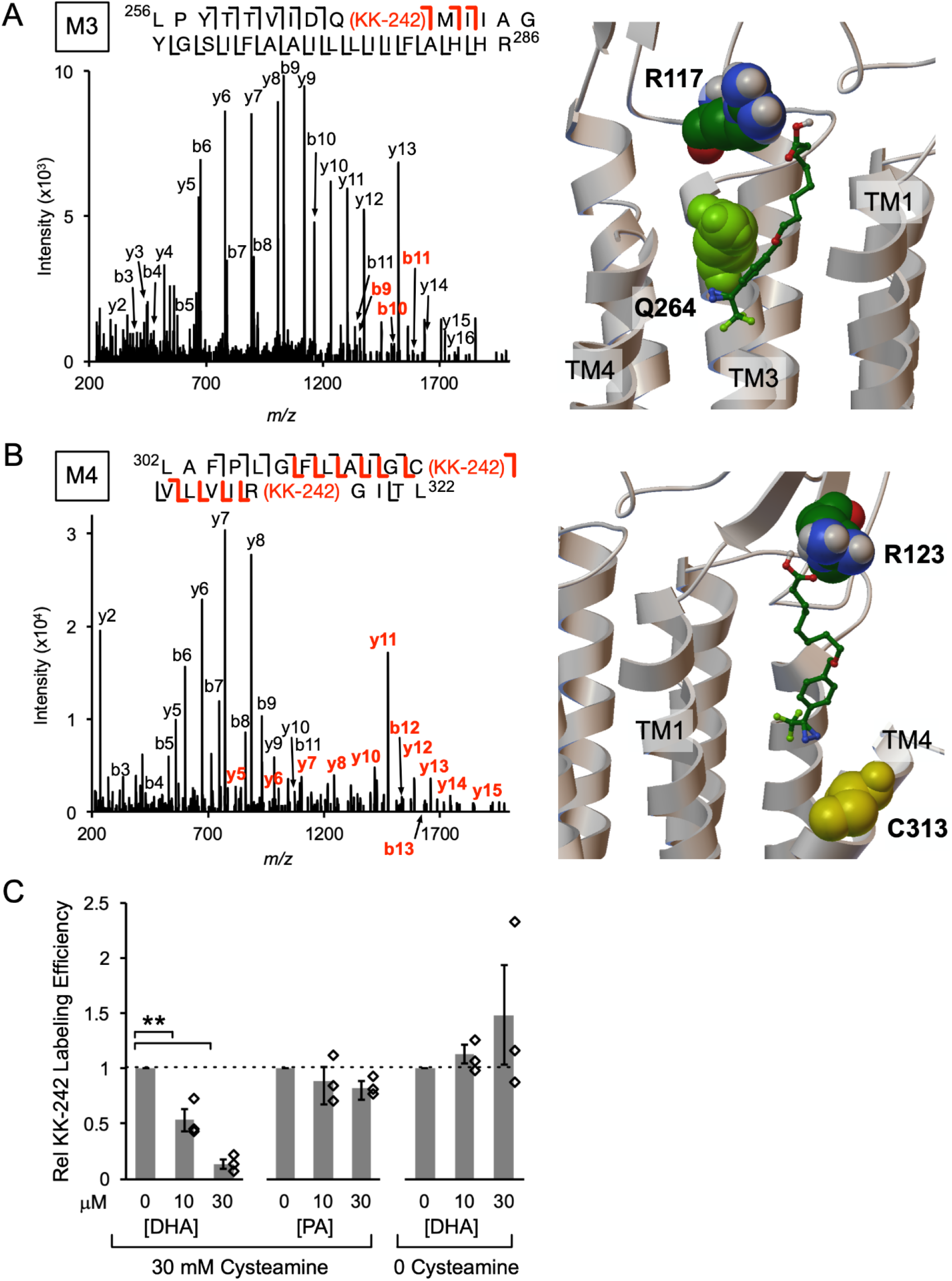
KK-242 photolabels two DHA binding sites in the outer TMD of ELIC. (A) *Left:* Fragmentation spectrum (MS2) of a M3 peptide photolabeled with KK-242. Red *b*-ions contain the KK-242 adduct mass, while black *b-* and *y*-ions do not. The photolabeled residue, Q264, is indicated with KK-242 in parenthesis. *Right:* ELIC structure showing a KK-242 binding pose from docking with the diazirine adjacent to Q264. Also shown is R117. (B) *Left:* MS2 spectrum of a M4 peptide photolabeled with KK-242. The fragment ions indicate labeling at C313 and R318. *Right:* ELIC structure showing a KK-242 binding pose from docking with the diazirine adjacent to C313. Also shown is R123. (C) Photolabeling efficiencies of M4 by 10 μM KK-242 in the presence of 10 and 30 μM DHA and PA. For DHA, labeling efficiencies were evaluated in the presence and absence of cysteamine. Labeling efficiencies in the presence of DHA or PA were normalized to the labeling efficiency in the absence of fatty acid (n=3, ±SEM, ** p<0.01).

**Figure 4—figure supplement 1:**
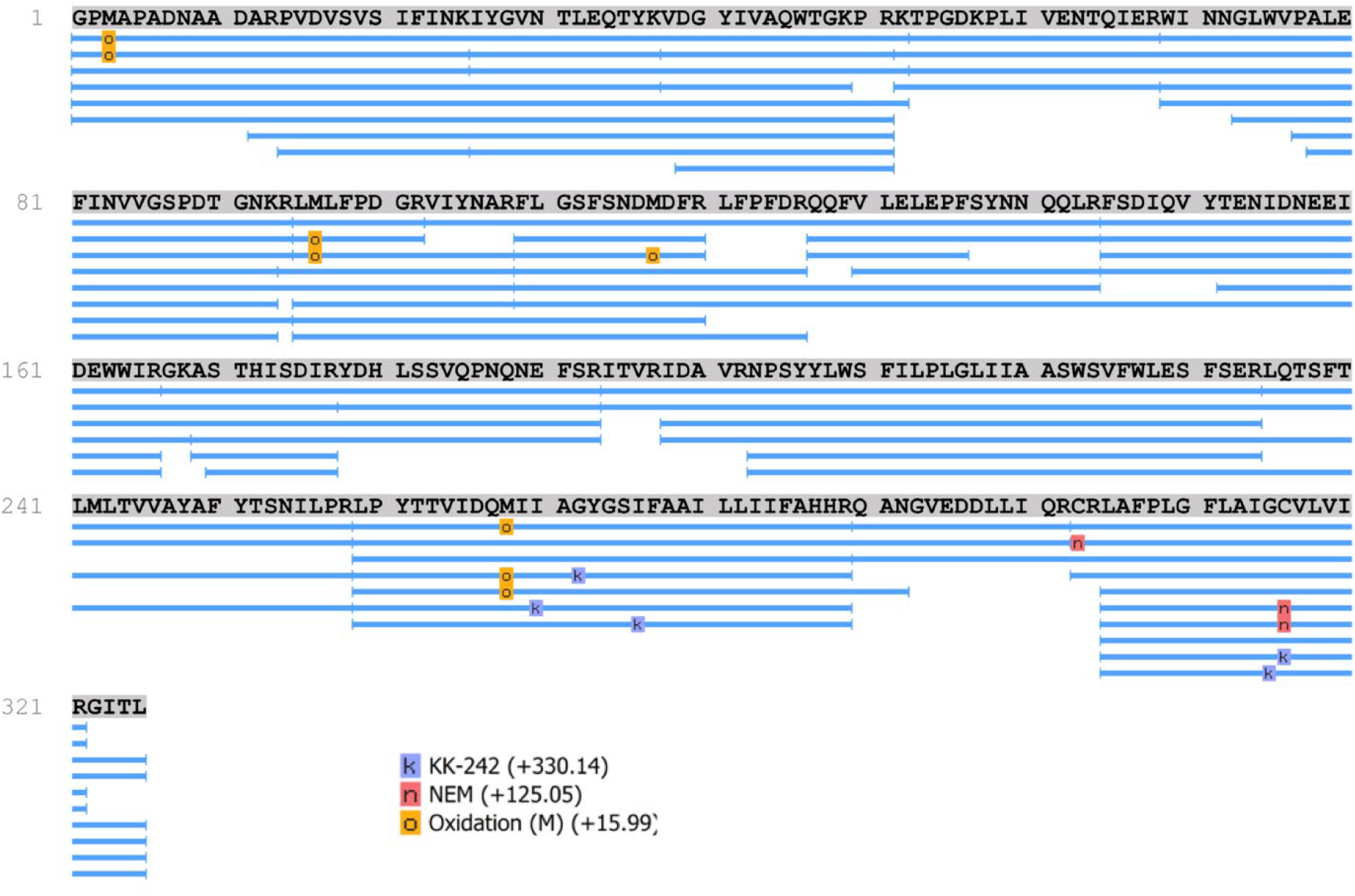
Peptide coverage map from a PEAKS search of the tryptic middle-down LC-MS data from ELIC photolabeled with 100 μM KK-242. Modifications used in the search include KK-242 (k), methionine oxidation (o), and cysteine alkylation with NEM (n). Peptide sequence coverage was 100%. The precursor mass accuracy was set at 20 ppm. KK-242 was identified in M3 and M4 peptides. The assigned residue modified by KK-242 in these peptides is not reliable due to the relatively low stringency set for fragment ion identification (0.1 Da accuracy). Manual analysis of these MS2 spectra confirmed the location of KK-242 when requiring a fragment ion mass accuracy of <10 ppm.

**Figure 4—figure supplement 2:**
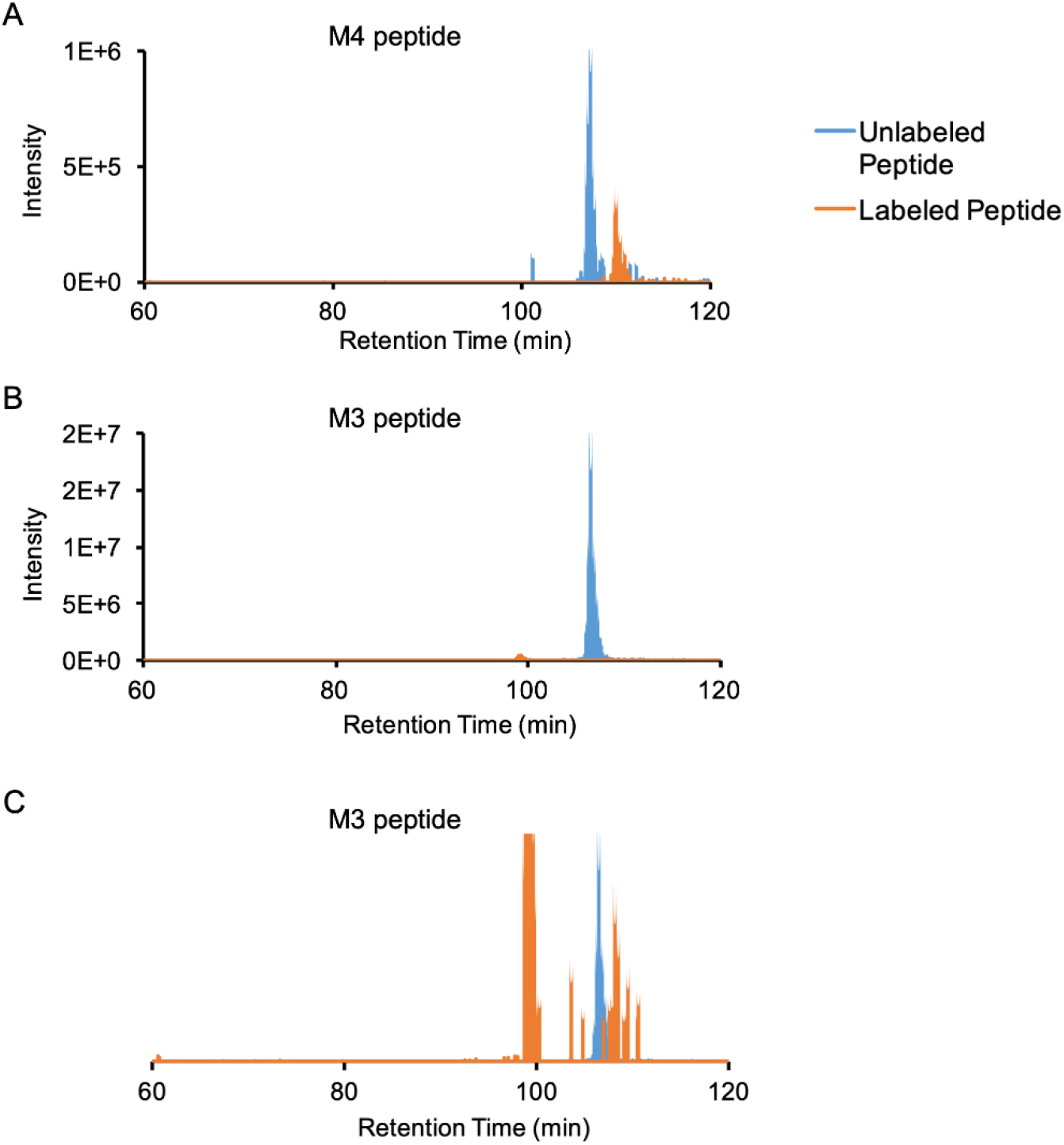
Extracted ion chromatograms of M4 and M3 unlabeled and labeled peptides. (A) Unlabeled (736.76-736.77 m/z) and labeled (839.48-839.50 m/z) extracted ion chromatograms of the M4 peptide (monoisotopic mass of labeled peptide = 939.491 m/z, z = 3+). The labeled peptide elutes ~3 min after the unlabeled peptide, and the estimated labeling efficiency is ~32%. (B) Unlabeled (865.23-865.24 m/z) and labeled (947.76-947.77 m/z) extracted ion chromatograms of the M3 peptide (monoisotopic mass of labeled peptide = 947.768 m/z, z = 4+). The estimated labeling efficiency is ~2.8%. (C) Same as (B) but with the labeled peptide ion chromatogram scaled up by 100x for better visualization: the labeled peptide elutes ~1.7 min after the unlabeled peptide. The peak at 100 min retention time comes from a different peptide with the same m/z but different charge state.

**Figure 4—figure supplement 3:**
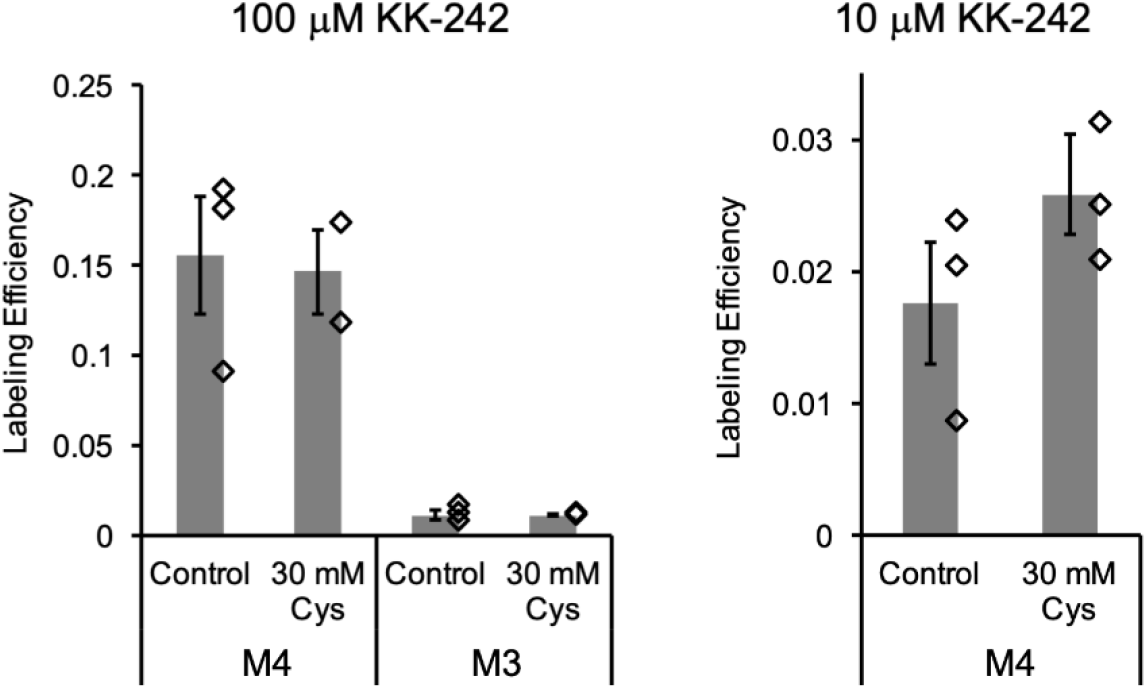
Photolabeling efficiencies determined from middle-down tryptic LC-MS analysis of M4 and M3 peptides in ELIC. ELIC was photolabeled with 100 μM (*left*) or 10 μM (*right*) KK-242 in the absence or presence of 30 mM cysteamine. Labeling efficiencies are reported as mean ± SEM (n=3, n=2 for 100 μM KK-242 + 30 mM cysteamine).

**Figure 4—figure supplement 4:**
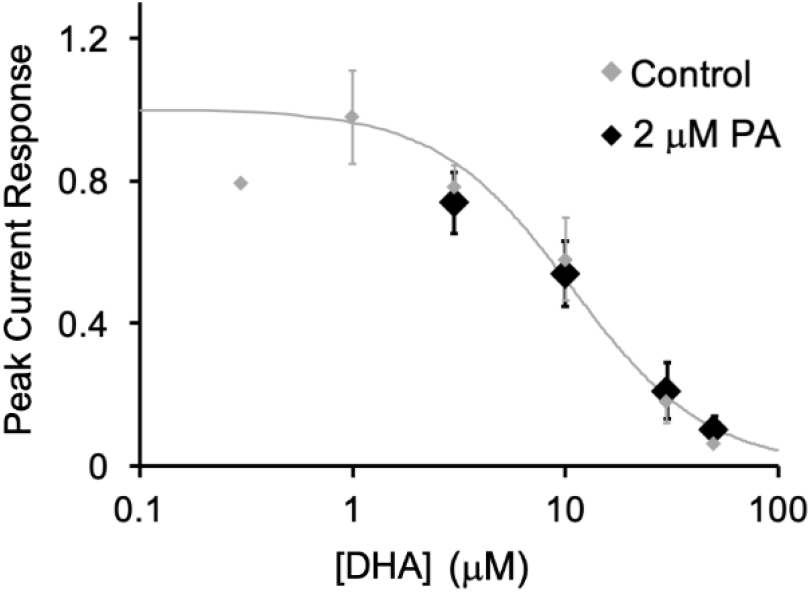
ELIC peak current responses to 30 mM cysteamine normalized to control (absence of DHA) as a function of DHA concentration. Black shows data in the presence of 2 μM PA (n=4, ±SEM). Gray shows data from Fig. 1B.

**Figure 4—table supplement 1:**
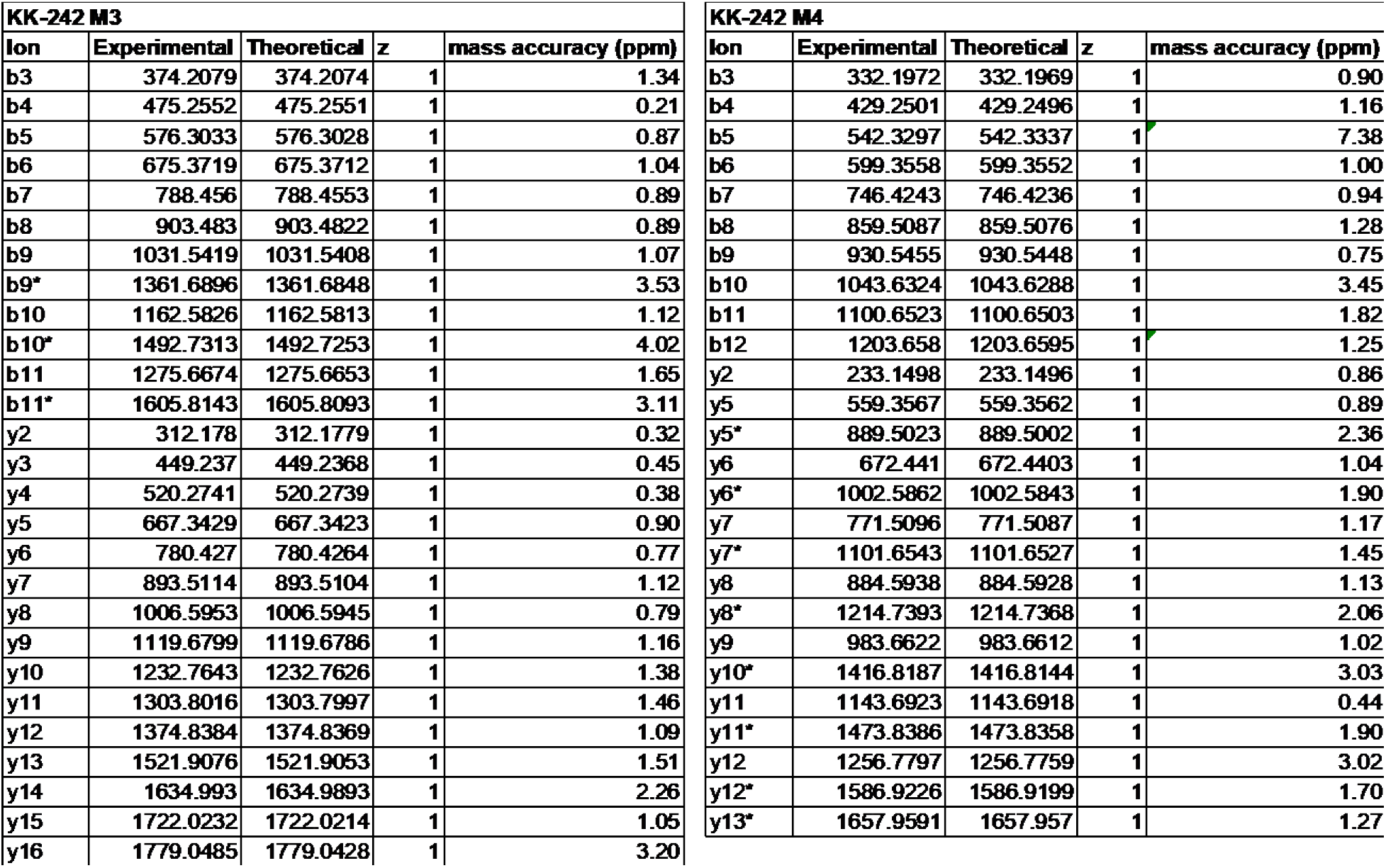
Experimental and theoretical m/z values from the fragmentation (MS2) spectra in Figure 2 for the KK-242 M3 and M4 peptides. * indicates *b* or *y* fragment ions containing the KK-242 mass. All fragment ions were confirmed to have a mass accuracy of < 10 ppm.

| KK-242 M3 |  |  |  |  | KK-242 M4 |  |  |  |  |
| --- | --- | --- | --- | --- | --- | --- | --- | --- | --- |
| Ion | Experimental | Theoretical | z | mass accuracy (ppm) | Ion | Experimental | Theoretical | z | mass accuracy (ppm) |
| b3 | 374.2079 | 374.2074 | 1 | 1.34 | b3 | 332.1972 | 332.1969 | 1 | 0.90 |
| b4 | 475.2552 | 475.2551 | 1 | 0.21 | b4 | 429.2501 | 429.2496 | 1 | 1.16 |
| b5 | 576.3033 | 576.3028 | 1 | 0.87 | b5 | 542.3297 | 542.3337 | 1 | 7.38 |
| b6 | 675.3719 | 675.3712 | 1 | 1.04 | b6 | 599.3558 | 599.3552 | 1 | 1.00 |
| b7 | 788.456 | 788.4553 | 1 | 0.89 | b7 | 746.4243 | 746.4236 | 1 | 0.94 |
| b8 | 903.483 | 903.4822 | 1 | 0.89 | b8 | 859.5087 | 859.5076 | 1 | 1.28 |
| b9 | 1031.5419 | 1031.5408 | 1 | 1.07 | b9 | 930.5455 | 930.5448 | 1 | 0.75 |
| b9* | 1361.6896 | 1361.6848 | 1 | 3.53 | b10 | 1043.6324 | 1043.6288 | 1 | 3.45 |
| b10 | 1162.5826 | 1162.5813 | 1 | 1.12 | b11 | 1100.6523 | 1100.6503 | 1 | 1.82 |
| b10* | 1492.7313 | 1492.7253 | 1 | 4.02 | b12 | 1203.658 | 1203.6595 | 1 | 1.25 |
| b11 | 1275.6674 | 1275.6653 | 1 | 1.65 | y2 | 233.1498 | 233.1496 | 1 | 0.86 |
| b11* | 1605.8143 | 1605.8093 | 1 | 3.11 | y5 | 559.3567 | 559.3562 | 1 | 0.89 |
| y2 | 312.178 | 312.1779 | 1 | 0.32 | y5* | 889.5023 | 889.5002 | 1 | 2.36 |
| y3 | 449.237 | 449.2368 | 1 | 0.45 | y6 | 672.441 | 672.4403 | 1 | 1.04 |
| y4 | 520.2741 | 520.2739 | 1 | 0.38 | y6* | 1002.5862 | 1002.5843 | 1 | 1.90 |
| y5 | 667.3429 | 667.3423 | 1 | 0.90 | y7 | 771.5096 | 771.5087 | 1 | 1.17 |
| y6 | 780.427 | 780.4264 | 1 | 0.77 | y7* | 1101.6543 | 1101.6527 | 1 | 1.45 |
| y7 | 893.5114 | 893.5104 | 1 | 1.12 | y8 | 884.5938 | 884.5928 | 1 | 1.13 |
| y8 | 1006.5953 | 1006.5945 | 1 | 0.79 | y8* | 1214.7393 | 1214.7368 | 1 | 2.06 |
| y9 | 1119.6799 | 1119.6786 | 1 | 1.16 | y9 | 983.6622 | 983.6612 | 1 | 1.02 |
| y10 | 1232.7643 | 1232.7626 | 1 | 1.38 | y10* | 1416.8187 | 1416.8144 | 1 | 3.03 |
| y11 | 1303.8016 | 1303.7997 | 1 | 1.46 | y11 | 1143.6923 | 1143.6918 | 1 | 0.44 |
| y12 | 1374.8384 | 1374.8369 | 1 | 1.09 | y11* | 1473.8386 | 1473.8358 | 1 | 1.90 |
| y13 | 1521.9076 | 1521.9053 | 1 | 1.51 | y12 | 1256.7797 | 1256.7759 | 1 | 3.02 |
| y14 | 1634.993 | 1634.9893 | 1 | 2.26 | y12* | 1586.9226 | 1586.9199 | 1 | 1.70 |
| y15 | 1722.0232 | 1722.0214 | 1 | 1.05 | y13* | 1657.9591 | 1657.957 | 1 | 1.27 |
| y16 | 1779.0485 | 1779.0428 | 1 | 3.20 |  |  |  |  |  |

To test whether KK-242 photolabeled sites are specific binding sites for DHA or PA, a competition experiment was performed: labeling efficiencies at 10 μM KK-242 were evaluated in the presence of 30 mM cysteamine with and without the addition of DHA or PA. At 10 μM KK-242, the labeling efficiency of M4 was 2.6% ± 0.5% (±SD, n=3) in the absence of DHA or PA (Figure 4—figure supplement 3); a photolabeled M3 peptide was not detectable at this lower concentration of KK-242. A decrease in labeling efficiency of M4 in the presence of DHA or PA would suggest that these fatty acids are competing for KK-242 at the photolabeled sites. While DHA caused a dose-dependent reduction in KK-242 labeling efficiency of M4, PA caused a small, statistically insignificant reduction (Figure 4C). This result indicates that DHA binds to KK-242 photolabeled sites with higher affinity than PA and is consistent with the inhibitory effects of DHA and PA on ELIC and the CGMD results. While we do not know the relative contribution of each site to M4 photolabeling, the large decrease in M4 photolabeling efficiency in the presence of DHA (~90% with 30 μM DHA) suggests that both photolabeled sites are occupied by DHA. Consistent with the specificity of these binding sites for DHA over PA, the potency of DHA inhibition of ELIC peak responses by giant liposome patch-clamping was not altered in the presence of 2 μM PA (Figure 4—figure supplement 4).

DHA inhibits ELIC peak responses but has minimal effect on the EC_50_ of cysteamine activation; this suggests that DHA inhibits ELIC by stabilizing an agonist-bound non-conducting state (34) (see Discussion). To test this possibility, we examined the effect of DHA on KK-242 labeling efficiency of M4 in the absence of cysteamine. Indeed, DHA did not reduce KK-242 labeling efficiency in the absence of cysteamine, in stark contrast to its effect in the presence of cysteamine (Figure 4C). Thus, DHA preferentially binds to KK-242 photolabeled sites when ELIC is in the agonist-bound state. It is worth noting that the photolabeling efficiency of 10 μM KK-242 in the presence of cysteamine was only modestly higher than in the absence of cysteamine (Figure 4—figure supplement 3). This is most likely because: 1) KK-242, being a weak inhibitor of ELIC, does not show the same state-dependence of binding as DHA and/or 2) the irreversible nature of photolabeling precludes detection of state-dependent binding. Nevertheless, the results clearly demonstrate that DHA preferentially binds to the photolabeled sites in an agonist-bound state.

Having identified two DHA binding sites on each face of M4 by photolabeling, we compared these results to the CGMD simulation data. We measured the contact probability of DHA with each amino acid in ELIC (Figure 5A, B, & D), to determine if DHA preferentially interacted with M1 or M3. While DHA has a non-zero contact probability with all lipid-facing transmembrane helices (*i.e*., M1, M3, and M4), residues on M3 have the highest cumulative contact probability (Figure 5—figure supplement 1). Interestingly, the residues with the highest contact probability between DHA and ELIC (A268, G271, and A275) are located on a single face of M3 (Figure 5D), namely the face oriented toward M4, suggesting a specific DHA binding site in the groove between M3 and M4. R117 and Q264 (Q264 is photolabeled by KK-242) are located on the same face of M3 and interact with DHA, especially the carboxylate headgroup throughout the simulation (Figure 5D, Figure 5—figure supplement 2). The remaining residues on the binding face of M3 (A268, G271, S272, and A275) interact extensively with the polyunsaturated tail. These results reinforce the photolabeling data and identify the M3-M4 intrasubunit groove as a specific binding site for DHA. While M3 is noted to have the highest averaged contact probability (7.8% in M3 vs 5.3% in M1 and 4.7% in M4), a lower, but non-trivial, amount of contact is noted between DHA and M1 (Figure 5A, Figure 5—figure supplement 1), consistent with KK-242 photolabeling of C313.

**Figure 5:**
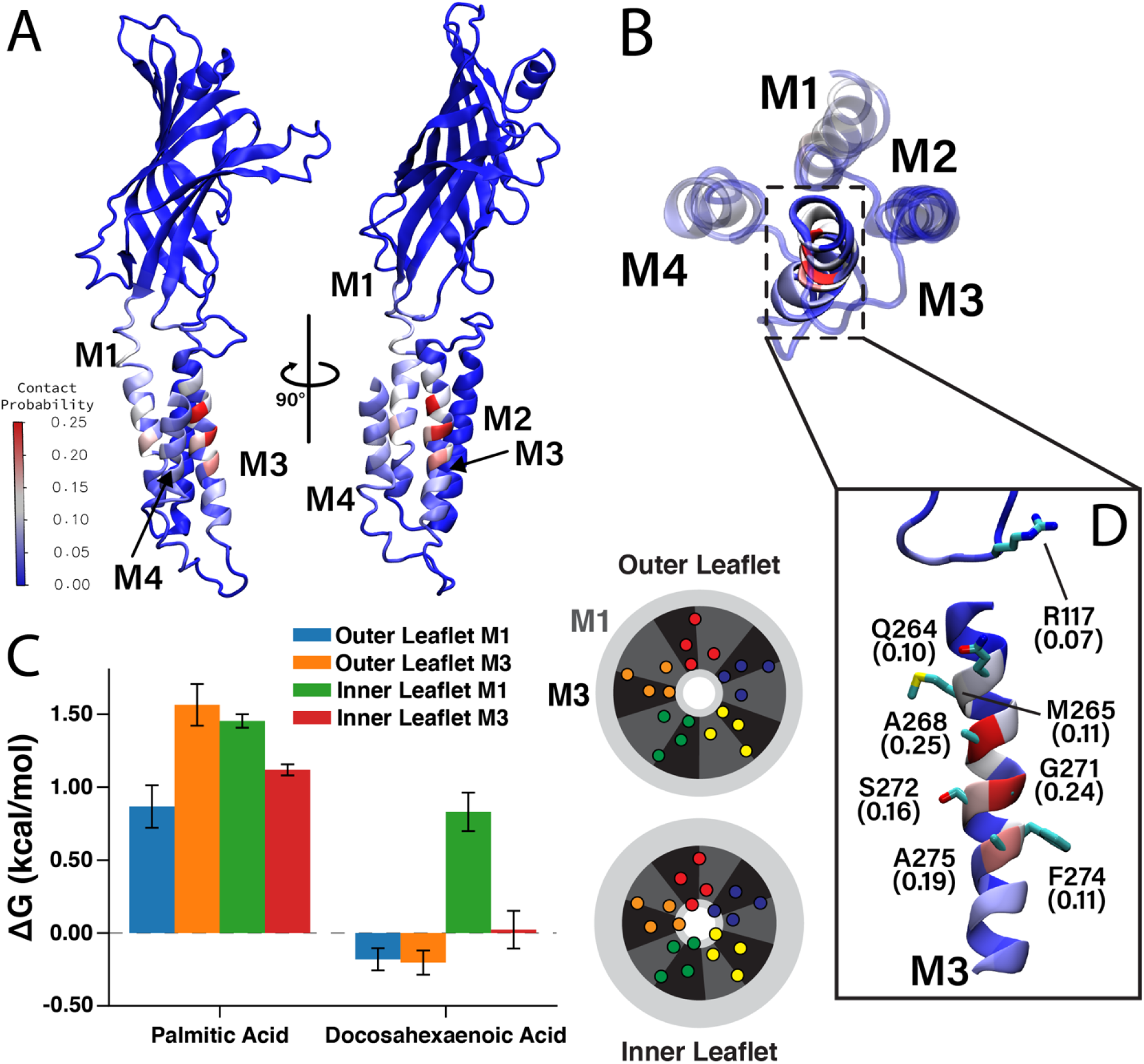
Favorable DHA binding modes from coarse-grained molecular dynamics. (A) An ELIC monomer is colored according to contact probabilities. The color bar to the left shows the contact probabilities values associated with the residue coloring. The same data is presented in (B) showing just the TMD as seen from the extracellular side. The M3 helix is shown in solid color, while M1, M2, and M4 are shown as transparent to highlight the DHA binding groove between M3 and M4. (C) Density threshold affinities for PA and DHA are shown (left) for the M1 and M3 sites in both the inner and outer leaflet (n=4, ±SEM). To the right is a cartoon representation of the boundaries for the M1 (dark gray) and M3 (black) sites with the colored helices representing the transmembrane domain. Each set of four same-colored helices represents one ELIC monomer. (D) The *β* 6-*β* 7 loop and M3 are colored according to contact probability. Residues in this binding site with the highest contact probabilities are highlighted by stick representation and the associated contact probabilities are shown.

To quantitatively compare the stability of DHA in each of these sites, we calculated a quantity analogous to a binding affinity, the density threshold affinity. The density threshold affinity ΔG (35) represents the relative free energy difference between a lipid species in the bulk membrane and in a specified, well-defined region near the protein, and is calculated using density ratios. Here, we define four sites per subunit, the “M1 site” (dashed sectors in Figure 2) in each of the two leaflets, and the “M3 site” (solid sectors in Figure 2) in each of the two leaflets. The site boundaries are inspired by the 2D density distributions in Figure 2, have five-fold symmetry, and encompass non-overlapping regions around each respective helix. The calculated values of ΔG for PA and DHA for each of the four sites are shown in Figure 5C. Consistent with radial enrichment results described earlier, DHA has higher affinity across all sites compared to PA. Indeed, PA has unfavorable interaction with all sites measured, having the lowest affinity for the outer leaflet M1 site (Figure 5C). Comparing affinity of DHA between different sites, there is a clear preference for outer leaflet sites versus inner leaflet sites, with both outer leaflet sites having favorable energy change and both inner leaflet sites having unfavorable energy change. This is consistent with the radial enhancement data and the fact that KK-242 was not observed to label inner leaflet sites. Although not statistically significant, there is slightly higher affinity of DHA for the outer membrane M3 site than the outer membrane M1 site (Figure 5C) in agreement with the contact probability analysis. Overall, the results from these analyses support the notion that DHA binds to outer leaflet intrasubunit sites.

**Figure 5—figure supplement 1:**
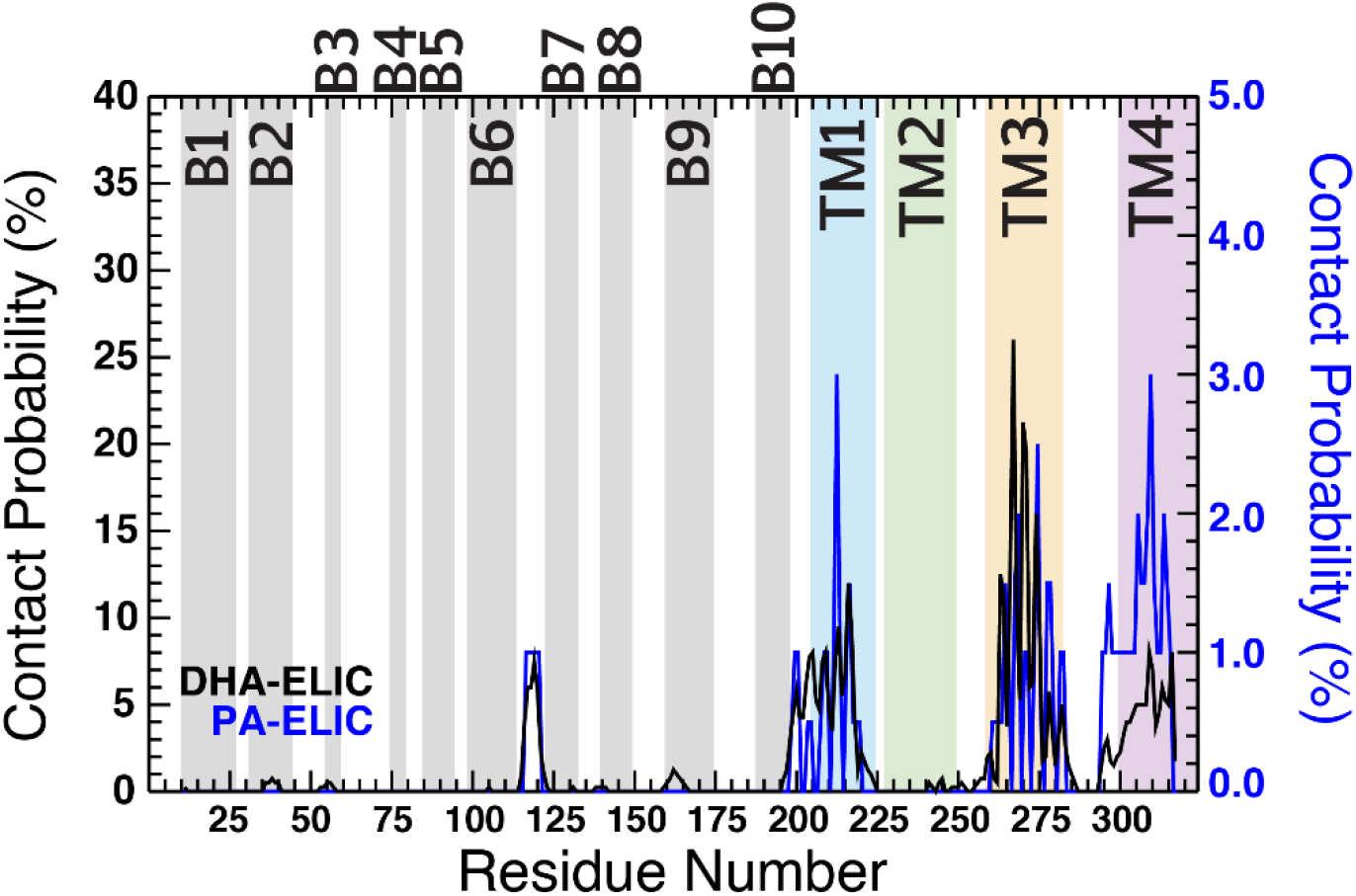
Contact probability is shown for PA (blue trace) and DHA (black trace) as a function of residue number. The scale for PA contact probability is located to the right of the plot and is lower in absolute value than the scale for DHA contact probability (left of the plot). The residue numbers making up the ECD β-sheets (β1-β10) are shown as gray fill in the background of the plot. The residue numbers comprising the TMD α-helices are shown as colored fill in the background of the plot (M1, blue; M2, green; M3, yellow; M4, purple). The contact probability represents an average across all five subunits for four simulation runs under each condition.

**Figure 5—figure supplement 2:**
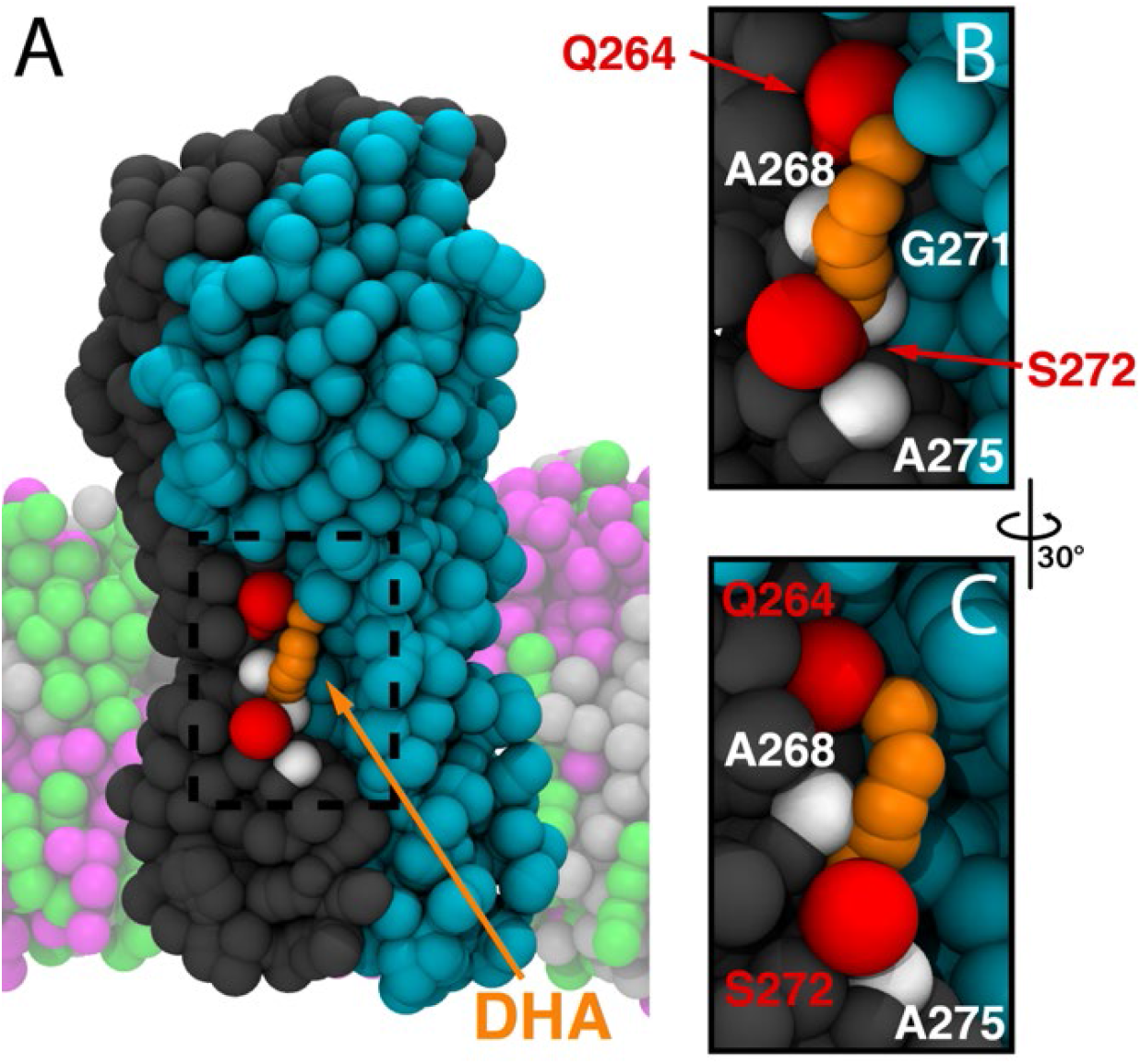
(A) Representative molecular image of DHA bound to ELIC from the CGMD simulations. Two complimentary ELIC monomers are shown in different colors (gray and teal) with the bound DHA shown as orange beads. Residues with high contact probability are shown as spheres with polar residues (Q264, S272) shown in red and apolar residues (A268, G271, A275) shown in white. In the coarse-grained membrane (translucent), POPC is pink, POPE is light gray, and POPG is green. The images in (B) and (C) focus on the dashed box in (A).

**Figure 5—figure supplement 3:**
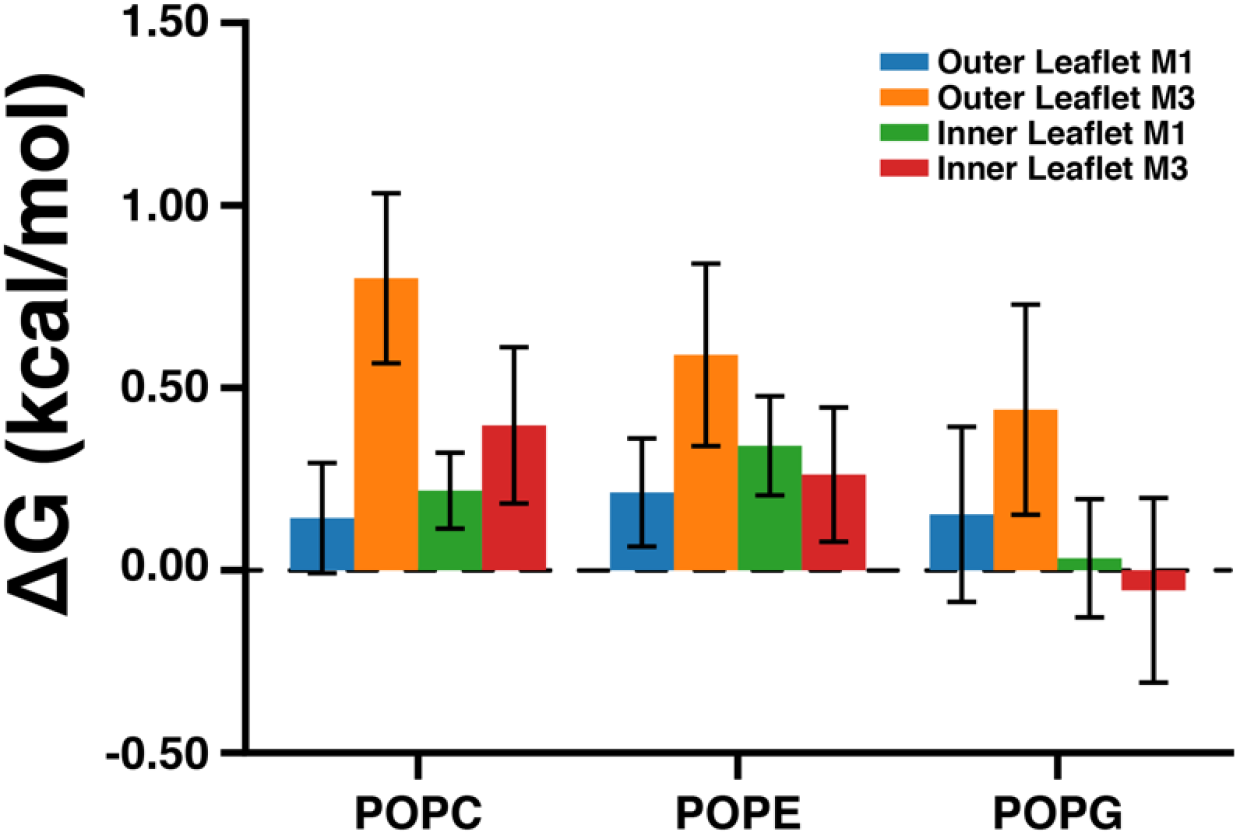
Density threshold affinities are plotted for POPC, POPE, and POPG from the CGMD simulations. This data represents an average across all five subunits and eight simulation runs (±SEM).

### DHA inhibits ELIC through one high affinity site

To investigate the functional significance of the M1 and M3 DHA binding sites, we modified each site with hexadecyl-methanethiosulfonate (hMTS); the hexadecyl group is identical to a PA alkyl tail. Based on the photolabeling, docking and CGMD results, we reasoned that the fatty acid headgroup is near R117 and R123 when occupying either binding site (Figure 4A and 4B, Figure 6A). Thus, we modified these positions with hMTS to mimic binding of a fatty acid. On a cysteine-less background (C300S/C313S mutant), we introduced R117C and R123C mutations. These triple mutants produced stable purified protein, and were incubated with 100 μM hMTS for 1 h and analyzed by intact protein MS. Both mutants were completely modified by one hMTS per subunit consistent with specific modification of the introduced cysteine (Figure 6A). We previously showed that a cluster of three arginines in the inner interfacial region (R286, R299, R301) of the ELIC TMD form charged interactions with anionic phospholipids, which influence channel agonist responses (Figure 6A) (24). Thus, we also generated cysteine mutations of these arginines; interestingly, these mutants were not modified by hMTS indicating decreased accessibility at this site (Figure 6A). This finding is consistent with CGMD results showing reduced binding of DHA and PA in the inner leaflet compared to the outer leaflet (Figure 2 and 5C). Moreover, analysis of the diacylated lipids species in the CGMD system (Figure 5—figure supplement 3) shows that POPG has a favorable affinity for this site, suggesting that higher occupancy of this site by POPG leads to the relative exclusion of fatty acids or hMTS.

**Figure 6:**
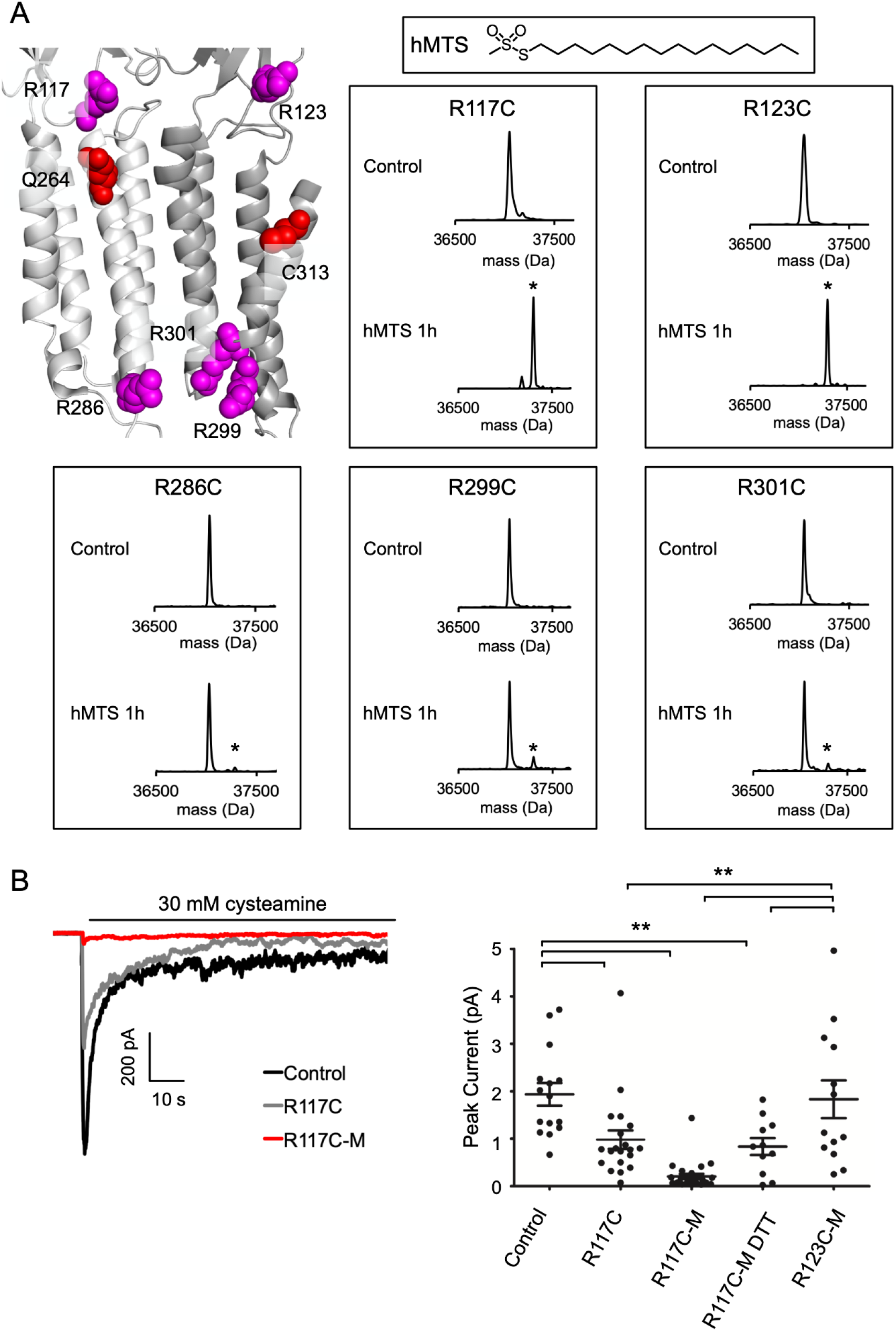
hMTS modification of R117C/C300S/C313S inhibits ELIC agonist response. (A) *Top left:* Structure of ELIC showing photolabeled residues (Q264 and C313), and arginines at the outer (R117, R123) and inner (R286, R299, R301) interfacial regions of the TMD. *Right:* Deconvoluted intact protein mass spectra of the indicated arginine to cysteine mutants on a C300S/C313S background. Mass spectra are shown prior to hMTS modification (control) and after 1h incubation with hMTS. The peak in all control spectra corresponds to the mass of unmodified ELIC mutant (37,048 Da), and a +257.5 Da shift indicates modification with one hMTS (marked by asterisk). (B) *Left:* Sample currents of control (C300S/C313S), R117C (C300S/C313S/R117C), and R117C-M from 2:1:1 POPC:POPE:POPG giant liposomes. *Right:* Average peak current from control (C300S/C313S), R117C (C300S/C313S/R117C), R117C-M, R117C-M treated with 2 mM DTT, and R123C-M in response to 30 mM cysteamine (n=12-26, ±SEM, ** p<0.01).

Having modified R117C/C300S/C313S and R123C/C300S/C313S with hMTS (henceforth called R117C-M and R123C-M), we re-purified the modified proteins by size exclusion chromatography (SEC) to confirm a monodisperse sample and reconstituted in giant liposomes for excised patch-clamp recordings. While this approach worked for R123C-M, R117C-M in DDM was partially aggregated by SEC. To enable liposome reconstitution of R117C-M, we first equilibrated unmodified R117C/C300S/C313S with DDM-destabilized liposomes, and then modified with hMTS such that R117C-M was maintained in a lipid environment (see Experimental Procedures). After this treatment, biobeads were added to complete liposome formation. Intact protein MS of R117C-M from this proteoliposome sample showed two main peaks corresponding to the modified mutant with and without a bound phospholipid (Figure 6—figure supplement 1A). This phospholipid adducted to a denatured ELIC subunit is likely due to the abundance of residual phospholipid in this sample. No unmodified mutant was detected; however, this spectrum had significantly more noise and adduction compared to ELIC samples in DDM. Based on this noise, we estimate that >90% of the R117C/C300S/C313S mutant was modified with hMTS but cannot rule out the presence of some unmodified protein. SDS-PAGE of protein from C300S/C313S and R117C-M proteoliposomes showed equal amounts of ELIC in both samples consistent with a successful reconstitution (Figure 6—figure supplement 1B).

Having produced liposomes with C300S/C313S, R117C-M or R123C-M, we examined channel responses to 30 mM cysteamine. Peak currents from excised patches of C300S/C313S and R123C-M were of similar magnitude, but peak currents of R117C-M were significantly smaller with many patches showing no current (Figure 6B). We also tested the R117C/C300S/C313S mutant (i.e. unmodified) and found that this mutation significantly reduced peak currents compared to C300S/C313S; however, R117C-M peak currents were still lower than R117C/C300S/C313S (significantly different based on student’s T-test but not ANOVA) (Figure 6B). To further verify that hMTS modification of R117C inhibits channel responses, we performed recordings from R117C-M proteoliposomes incubated in 2 mM DTT, which may remove the disulfide-mediated hMTS modification. R117C-M peak currents in the presence of 2 mM DTT were noticeably larger than in the absence of DTT (significantly different based on student’s T-test but not ANOVA) (Figure 6B). While there is substantial variability in peak currents between patches, the results suggest that hMTS modification of R117C inhibits channel responses, mimicking the effect of fatty acids. The C300S/C313S mutant produced slower current decay and larger steady state currents compared to WT, which is consistent with previous reports of the effect of mutations at these residues (36). However, cysteamine responses for R117C/C300S/C313S, R117C-M and R123C-M did not show significantly different rates of current decay or relative steady state current compared to C300S/C313S (Figure 6—figure supplement 2A). Moreover, all four showed similar relative reduction of peak current with preapplication of 30 μM DHA (Figure 6—figure supplement 2B).

**Figure 6—figure supplement 1:**
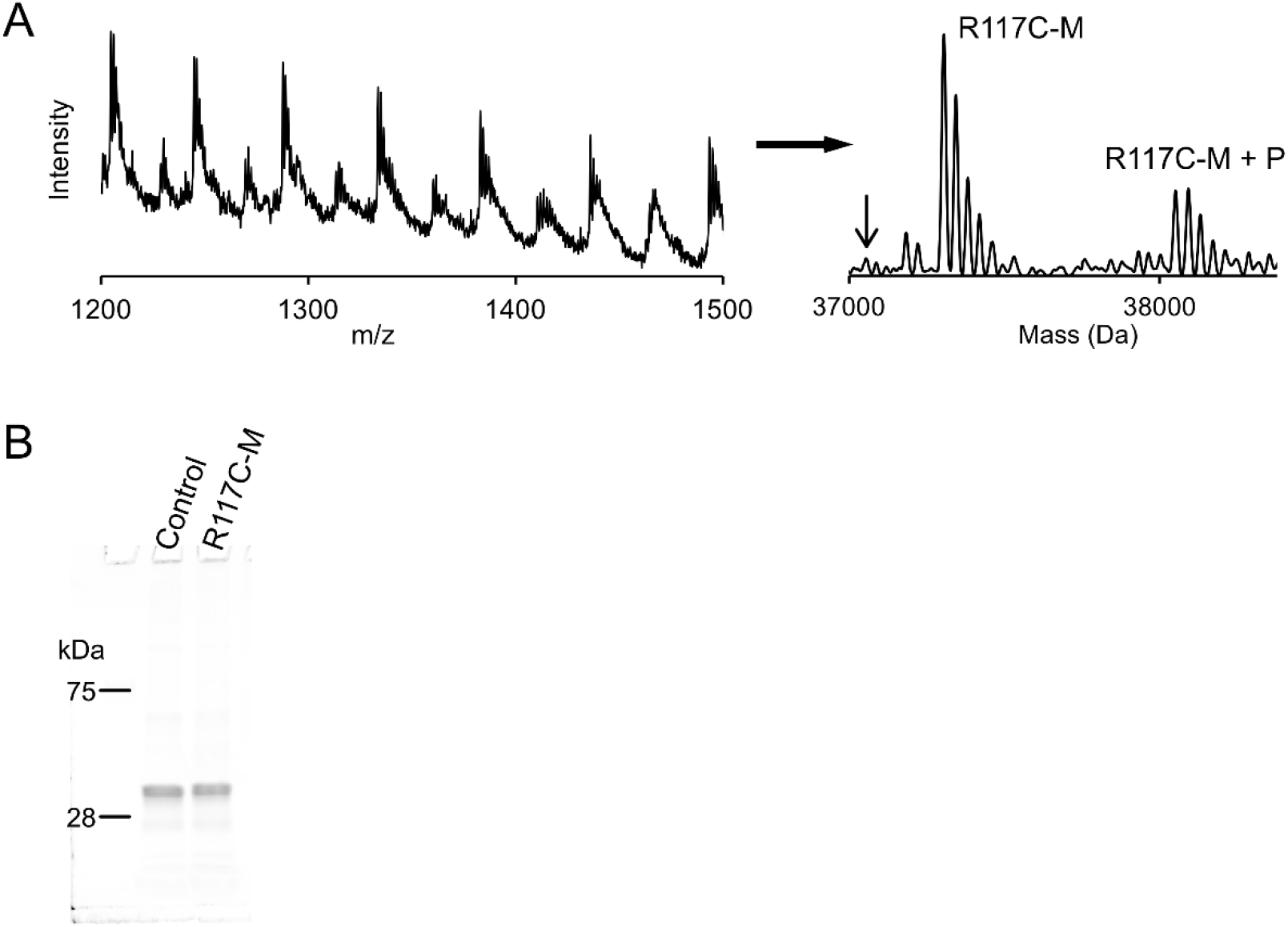
(A) *Left:* Full intact protein mass spectrum of R117C-M. *Right:* Deconvoluted mass spectrum of R117C-M with curved baseline subtraction to eliminate noise. The peaks labeled R117C-M correspond to the hMTS modified R117C/C300S/C313S subunit (37,305 Da) with 4-5 unknown small molecule adducts (+37-39 Da). The peaks labeled R117C-M + P are shifted by +746 Da corresponding to R117C-M with a bound POPG phospholipid. Arrow indicates the mass of unlabeled R117C/C300S/C313S (37,058 Da). (B) SDS-PAGE showing the ELIC protein subunit (~37 kDa) extracted from control (C300S/C313S) and R117C-M proteoliposomes.

**Figure 6—figure supplement 2:**
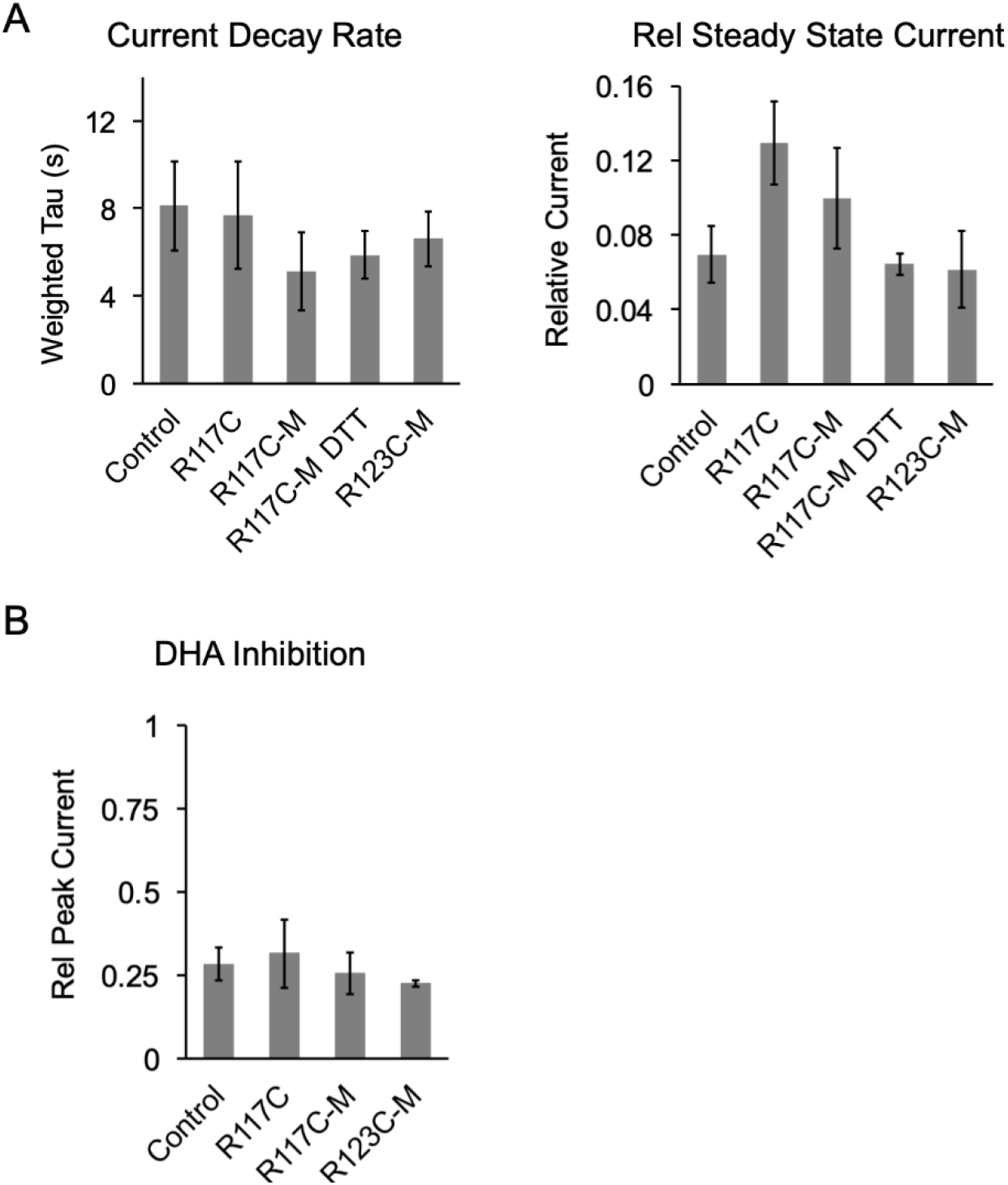
(A) From the current responses to 30 mM cysteamine from Fig. 5B, graphs show the weighted tau of current decay and steady state current normalized to peak (n=5-13, ±SEM). R117C is R117C/C300S/C313S. (B) Magnitude of DHA inhibition quantified as peak current after 3 min pre-application with 30 μM DHA normalized to peak current in the absence of DHA (n=5-9, ±SEM).

## DISCUSSION

Using a novel fatty acid photolabeling reagent and CGMD, we identified two fatty acid binding sites in ELIC, which are occupied by the PUFA, DHA, with much higher affinity than the saturated fatty acid, PA. The striking agreement between photolabeling and simulation results provides a compelling molecular description of the selectivity of fatty acid interactions with a pLGIC. Along with site-directed hMTS modification to interrogate the functional significance of these sites, the results strongly indicate that DHA inhibits ELIC through a single site, an intrasubunit groove between the top of M3 and M4. This site is analogous to the DHA binding site in a GLIC crystal structure (3). That this fatty acid binding site is present in both ELIC and GLIC suggests that it may also be conserved in eukaryotic pLGICs. Indeed, the equivalent site in the β3 subunit of the α1β3 GABA_A_R was found to be a neurosteroid binding site that mediates neurosteroid inhibition of GABA_A_R responses (37). Thus, this site may be a conserved lipid binding site through which fatty acids and neurosteroids inhibit pLGICs.

DHA binding to KK-242 photolabeled sites was specific for the agonist-bound state. This result supports a direct binding mechanism in which DHA allosterically inhibits ELIC responses by binding to and stabilizing an agonist-bound non-conducting state. Our patch-clamp data show that fatty acids inhibit ELIC peak responses when applied prior to agonist similar to previous reports in the GABA_A_R and nAchR. This is in contrast to the effect of DHA described in GLIC, in which DHA was co-applied with agonist resulting in no change in peak responses but faster and greater current decay (3). This apparent increase in the rate and extent of current decay was interpreted as fatty acid stabilization of a desensitized state. However, a different hypothesis was recently proposed by Gielen and Corringer for understanding the 1) increase in desensitization rate with fatty acid co-application and 2) decrease in peak response with fatty acid pre-application; both of these findings can be predicted by a model in which a pre-active state is stabilized (34). Our results with ELIC show that pre-application of DHA not only inhibits peak current, but also results in faster and more profound current decay. We have repeated the simulations performed by Gielen and Corringer using the same gating model (34), and found that the qualitative effects of fatty acid inhibition observed in ELIC (i.e. both a decrease in peak current and increase in the rate/extent of current decay with fatty acid pre-application) can be produced by stabilizing either a pre-active (Figure 6—figure supplement 3) or desensitized state (Figure 6—figure supplement 4). The model predicts that stabilization of a pre-active or desensitized state, to the extent that peak responses are completed inhibited, will lead to most channels occupying these respective states at steady-state (Figure 6—figure supplement 3C and 4C). Thus, more structures of pLGICs with fatty acids in the presence of agonist may reveal whether fatty acids stabilize pre-active or desensitized conformations of the channel.

**Figure 6—figure supplement 3:**
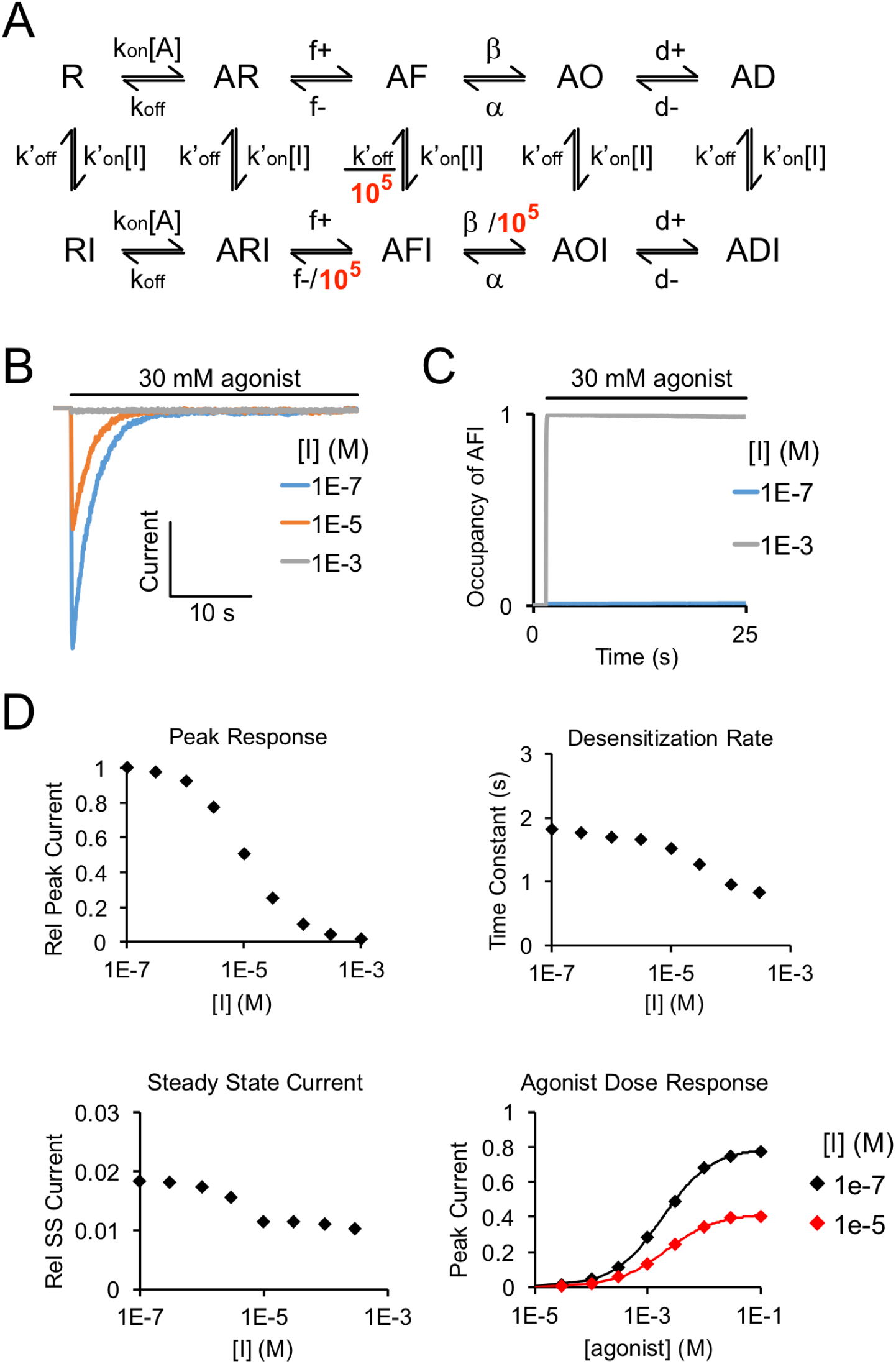
Simulated agonist responses demonstrating the effect of stabilizing an inhibitor (I)-bound pre-active state (AFI). (A) Model used for simulating agonist responses replicated from Gielen and Corringer (37) with adjustments made to the rate constants to approximate ELIC responses. R = resting, F = flipped (pre-active), O = open, D = desensitized. States with I represent inhibitor-bound states (e.g. DHA). k_on_ = 4,400 M^-1^s^-1^, k_off_ = 80 s^-1^, f^+^ = 50 s^-1^, f^-^ = 5000 s^-1^□ = 10,000 s^-1^, α = 100 s^-1^, d^+^ = 0.7 s^-1^, d^-^ = 0.01 s^-1^, k’_on_ = 10,000 M^-1^s^-1^, k’_off_ = 0.1 s^-1^. To reproduce the effect of DHA inhibition of ELIC responses, rate constants leaving AFI were decreased by 10^5^-fold. (B) Simulated currents were generated by applying 30 mM agonist and pre-applying inhibitor. (C) Simulation shows the probability of occupying the inhibitor-bound pre-active state (AFI) with 30 mM agonist, and high (1 mM) and low (0.1 μM) concentrations of inhibitor. High concentrations of inhibitor drive the channel entirely into the AFI state at steady state. (D) From the simulated responses, graphs show decrease in peak response, increase in the rate of current decay, and decrease in steady state current with increasing concentration of inhibitor. The inhibitor leads to a small left-shift in the agonist dose response curve (EC_50_ = 2.1 mM for [I] = 1e-7 M, EC_50_ = 1.8 mM for [I] = 1e-5 M).

**Figure 6—figure supplement 4:**
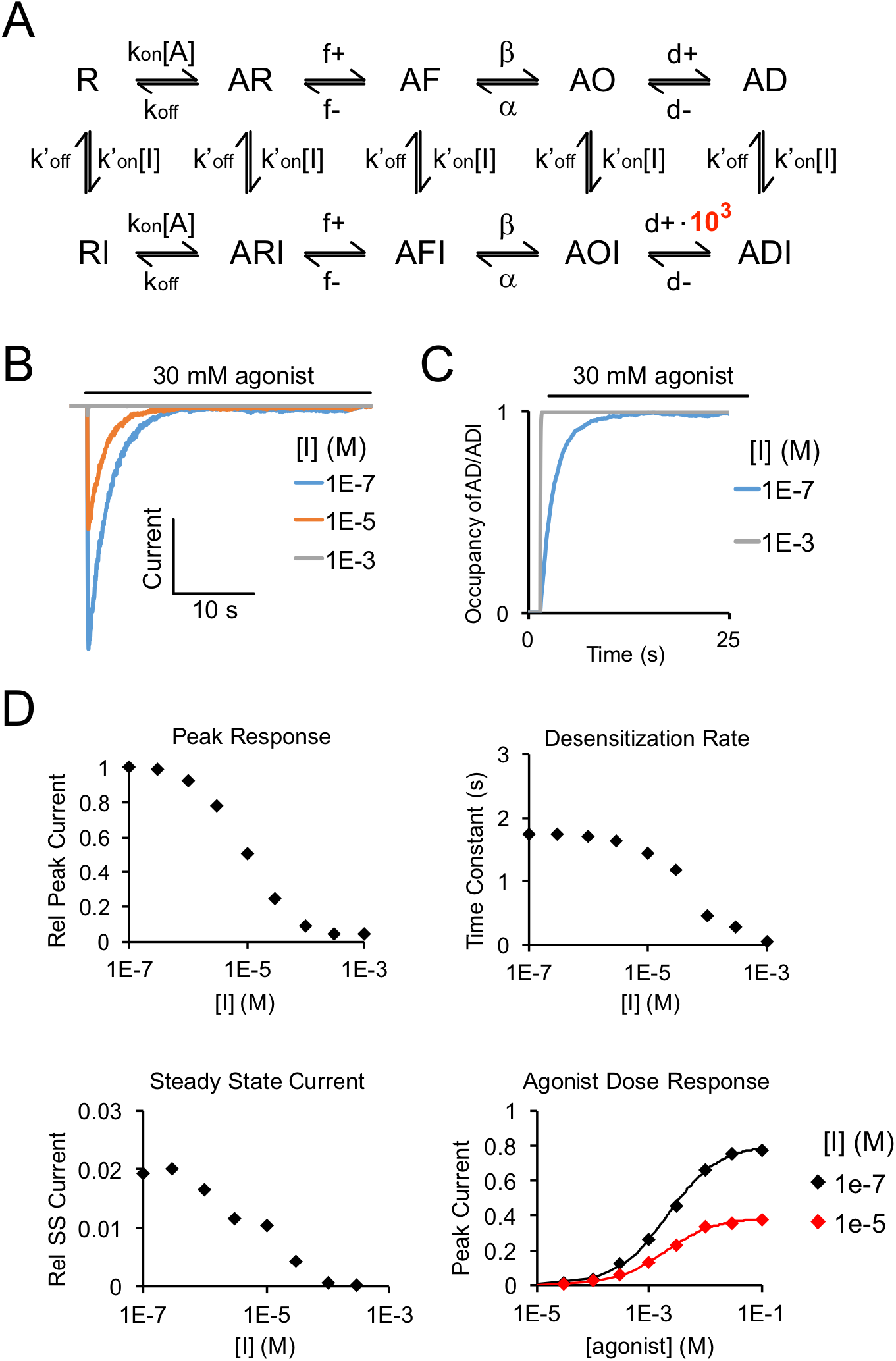
Simulated agonist responses demonstrating the effect of stabilizing an inhibitor (I)-bound desensitized state (ADI). (A) Same model and rate constants as Figure 5 Supplement 3, except to reproduce the effect of DHA inhibition of ELIC responses, the rate (d^+^) entering ADI (inhibitor-bound desensitized state) was increased by 10^3^-fold. (B) Simulated currents were generated by applying 30 mM agonist and pre-applying inhibitor. (C) Simulation shows the probability of occupying the inhibitor-bound desensitized state (ADI) with 30 mM agonist, and high and low concentrations of inhibitor. High concentrations of inhibitor drive the channel entirely into the ADI state at steady state. (D) Simulated responses show inhibition of peak response, increase in the rate of current decay, and lower steady state current with increasing concentration of inhibitor. The inhibitor leads to a small left-shift in the agonist dose response curve (EC_50_ = 2.1 mM for [I] = 1e-7 M, EC_50_ = 1.9 mM for [I] = 1e-5 M).

By introducing a hexadecyl group to fatty acid binding sites using hMTS, we sought to mimic occupancy of these sites by PA. Alkyl-MTS modification has been previously used to examine the effects of phospholipids at specific sites in inward rectifying potassium channels (38,39). In this study, we combined hMTS modification with intact protein MS to measure modification efficiency. This proved useful in showing that hMTS has limited access to a cluster of arginine residues in the inner TMD of ELIC, shown previously to be a binding site for the anionic phospholipid, phosphatidylglycerol (PG) (23,24,40). Since ELIC in DDM co-purifies with PG from *E. coli* membranes (24), it is plausible that hMTS cannot access these sites due to the presence of tightly bound PG. KK-242 may have not photolabeled sites in the inner TMD for the same reason. Indeed, the coarse-grained simulations with membranes containing 25% POPG showed significant reduction in the binding of DHA to the inner TMD of ELIC due to enrichment of POPG at this site. It is interesting to note that fatty acids are thought to modulate Kv channels by interacting with positively charged residues also in the extracellular side of the voltage-sensor domain (14,41,42). Thus, fatty acids may be excluded from sites in the inner TMD of many ion channels due to the presence of anionic phospholipids, which are localized in the inner leaflet of the cell membrane and bind with relatively high affinity to sites in the inner TMD (38,43–46).

It is reasonable to assume that the carboxylate anion or carboxylic acid of DHA interacts with interfacial arginines such as R117 in ELIC. Indeed, the carboxylic acid of DHA interacts with an arginine in the GLIC DHA structure (3). However, we found that DHA methyl ester inhibited ELIC with similar potency and efficacy as DHA meaning that an anionic carboxylate headgroup is not required for inhibition. Consistent with this finding, the R117C/C301S/C314S mutant showed similar sensitivity to DHA inhibition compared to C301S/C314S, and coarse-grained simulations of PA versus DHA clearly show that the polyunsaturated tail is the primary determinant of binding affinity. Thus, while CGMD and docking results suggest that the carboxylate headgroup of DHA is near R117 when bound to the M3/M4 site, a salt bridge is not a major determinant for DHA binding or inhibition in ELIC. This is consistent with a prior report where both DHA and DHA methyl ester were found to modulate hippocampal excitability through the GABA_A_R (10) suggesting that this mechanism of PUFA inhibition is conserved in mammalian pLGICs.

The strong inhibitory effect of alkylation of R117C is similar to the maximal effect of DHA or DHA methyl ester, but different from PA which has a much weaker effect. A likely explanation for this discrepancy is that PA has lower occupancy of this site compared to DHA or a covalently attached hMTS, resulting in a lower apparent efficacy. Indeed, PA was much less effective in reducing KK-242 photolabeling compared to DHA and was depleted around ELIC in the coarse-grained simulations. Thus, the difference in inhibition between DHA and PA appears to be entirely related to different binding affinities for this site. If a hexadecyl group resembling PA is covalently introduced to this site, it is also effective at inhibiting ELIC.

If hMTS modification of R117C occupies a DHA binding site, it may occlude the inhibitory effect of DHA; this was not the case with R117C-M. Since the small currents observed in R117C-M (~10% of control) remained sensitive to DHA, it is possible that these currents arise from ~10% of R117C/C300S/C313S that was not modified. This unmodified protein may have been missed by intact protein MS due to the high noise in the spectrum. We also cannot exclude the possibilities that: 1) DHA can still partially displace a covalently attached hMTS from this site, or 2) DHA also acts through an entirely different mechanism. Regardless of what accounts for these small DHA-sensitive currents, the strong inhibitory effect of R117C-M and the reversal of this effect by DTT support the hypothesis that this site mediates fatty acid inhibition of ELIC.

The photolabeling results with ELIC indicate that KK-242 is a useful fatty acid analogue photolabeling reagent. KK-242, which features a TPD, is advantageous compared to pacFA, which features an aliphatic diazirine, for the identification of fatty acid binding proteins and binding sites. This is likely because the TPD can label any amino acid including aliphatic and aromatic side chains, which usually line the binding pockets of fatty acid alkyl tails (47). This limitation of pacFA (i.e. preferential labeling of nucleophilic residues by aliphatic diazirines) likely applies to other lipid analogue photolabeling reagents that feature an aliphatic diazirine in the hydrophobic regions of the molecule (e.g. PhotoClick-Cholesterol, pacFA ceramide) potentially reducing the utility of these reagents for identifying lipid binding proteins (17,48,49).

In conclusion, DHA inhibits ELIC through specific binding to an M3/M4 intrasubunit site in the agonist-bound state.

## EXPERIMENTAL PROCEDURES

### Synthesis of KK-242

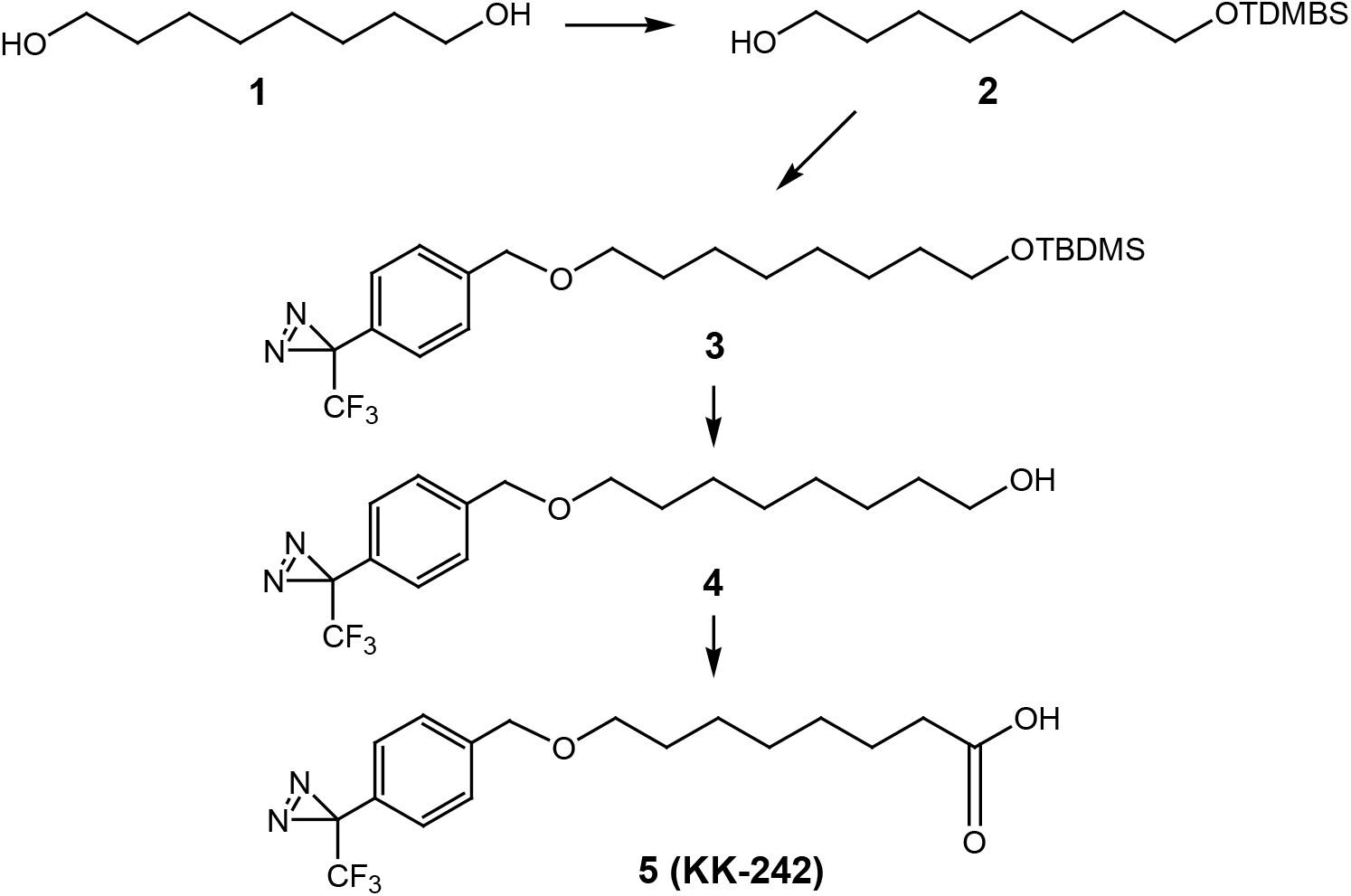

#### 8-(*tert*-Butyl-dimethyl-silanyoxy-1-ol (2)

1,8-Octanediol (**1**,730 mg, 5mmol), imidazole (408 mg, 6 mmol) and *t*-butyldimethylsilyl chloride (750 mg, 5 mmol) were stirred in DMF (8 mL) for 15 hours. Saturated aqueous NH_4_Cl was added to the reaction and the product was extracted into ethyl acetate. The combined ethyl acetate extracts were washed with brine, dried over anhydrous Na_2_SO_4_, filtered and the solvent removed under reduced pressure on a rotary evaporator. Flash column chromatography on silica gel yielded purified product **2** (630 mg, 48.4%). ^1^H NMR (400 MHz, CDCl_3_) δ 3.68-3.54 (m, 4H), 1.65-1.20 (m, 12H), 0.90 (s, 9H), 0.05 (s, 6H).

#### 3-[4-(8-*tert*-Butyl-dimethyl-silanyloxy)-octyloxymethylphenyl]-3-trifluoromethyl-3*H*-diazirine (3)

Sodium hydride in mineral oil (400 mg, 10 mmol), 3-(4-iodomethyl-phenyl)-3-trifluoromethyl-3*H*-diazirine (300 mg, 0.92 mmol) and compound **2** (200 mg, 0.77 mmol) in THF (15 mL) were heated at reflux for 2 hours (it is critical to stop the reaction after 2-hours to prevent product decomposition). The reaction mixture was cooled to 0 °C and the excess sodium hydride were carefully quenched by adding cold water. Additional water (50 mL) was added and the product was extracted into ethyl acetate (40 mL x 3). The combined organic extracts were washed with brine, dried over anhydrous Na_2_SO_4_, filtered and the solvent removed under reduced pressure on a rotary evaporator to give the compound **3** as a yellow oil which was passed through a short column of silica gel to remove less polar impurities. The column was first eluted with hexanes followed by 5% ethyl acetate in hexanes to give the product **3**, which was sufficiently pure to be converted to compound **4**.

#### 8-[4-(3-Trifluoromethyl-3*H*-diazirin-3-yl)benzyloxy-octan-1-ol (4)

Partially purified compound **4** was dissolved in methanol (5 mL) and a freshly prepared 5% dry-HCl-methanol solution made by adding acetyl chloride to methanol (3 mL) was added and the reaction was stirred for 1 hour. The reaction was made basic by adding aqueous saturated NaHCO_3_, and the product was extracted into dichloromethane (40 mL x 3). The combined organic extracts were washed with brine, dried over anhydrous Na_2_SO_4_, filtered and the solvent removed under reduced pressure on a rotary evaporator to give an oil which was purified by column chromatography (silica gel eluted with 15-25% ethyl acetate in hexanes) to give compound **4** (90 mg, 34%). ^1^H NMR (400 MHz, CDCl_3_) δ 7.37 (d, 2H, *J* = 7.2 Hz), 7.18 (d, 2H, *J* = 7.1Hz), 4.50 (s, 2H), 3.63 (t, 2H, *J* = 6.2 Hz), 3.46 (t, 2H, *J* = 6.2 Hz), 1.70-1.20 (m, 12H). ^13^C NMR (100 MHz, CDCl_3_) δ 140.62, 128.16, 127.76 (2 x C), 126.50 (2 xC), 122.14 (q, *J* = 275 Hz), 72.01, 70.77, 62.95, 32.72, 29.67, 29.38, 29.33, 26.08, 25.66.

#### 8-[4-(3-Trifluoromethyl-3*H*-diazirin-3-yl)benzyloxy-octanoic acid (5, KK-242)

Jones reagent (a few drops) was added to compound **4** (30 mg, 0.087 mmol) in acetone (5 mL) and the reaction was stirred for 2 hours. Excess Jones reagent was consumed by adding isopropyl alcohol (a few drops). The resulting blue solution was diluted with water (40 mL) and the product was extracted into ethyl acetate (30 mL x 3). The combined ethyl acetate extracts were washed with brine, dried over anhydrous Na_2_SO_4_, filtered and the solvent removed under reduced pressure on a rotary evaporator to give an oil which was purified by flash column chromatography (silica gel eluted with 20-50% ethyl acetate in hexanes) to give compound **5 (KK-242**, 24 mg, 77%). ^1^H NMR (400 MHz, CDCl_3_) δ 7.37 (d, 2H, *J* = 7.1 Hz), 7.19 (d, 2H, *J* = 7.1Hz), 4.51 (s, 2H), 3.46 (t, 2H, *J* = 6.2 Hz), 2.36 (t, 2H, *J* = 6.1 Hz), 1.70-1.200 (m, 10 H); ^13^C NMR (100 MHz, CDCl_3_) δ 179.69, 140.56, 128.15, 127.78 (2 x C), 126.52 (2 x C), 122.14 (q, *J* = 275 Hz), 72.02, 70.68, 33.92, 29.60, 29.01, 28.93, 25.94, 24.57.

### Expression, purification and mutagenesis of ELIC

ELIC was expressed and purified as previously described (24), yielding purified ELIC in 10 mM Tris pH 7.5, 100 mM NaCl, and 0.02% DDM (Buffer A). Site-directed mutagenesis was performed by a standard QuikChange approach and all mutations were confirmed by sequencing (Genewiz).

### ELIC giant liposome formation and excised patch-clamp recordings

ELIC WT or mutants were reconstituted in giant liposomes as previously described (24). For each liposome preparation, 0.5 mg of ELIC protein was reconstituted in 5 mg of 2:1:1 POPC:POPE:POPG liposomes, and the entire procedure as well as patch-clamp recordings was performed using 10 mM MOPS pH 7 and 150 mM NaCl (MOPS buffer). Giant liposomes were formed by dehydrating 10 μl of proteoliposomes on a glass coverslip in a desiccator for 1 h at RT followed by rehydration with 100 μl MOPS buffer overnight at 4 °C and 2 h at RT the following day. Giant liposomes were suspended by pipetting and then applied to the recording bath in MOPS buffer.

Excised patch-clamp recordings of ELIC in giant liposomes were performed as previously described (24). Bath and pipette solution contained MOPS buffer and 0.5 mM BaCl_2_. Fatty acid-containing solutions were prepared from DMSO stocks of 30 mM for each fatty acid. Thus, the final concentration of DMSO was 0.1% or less in all fatty acid-containing solutions, and 0.1% DMSO was added to control solutions to maintain consistency. Treatment of R117C-M proteoliposomes with DTT was performed by incubating the proteoliposomes in 2 mM DTT, and including 2 mM DTT in control and 30 mM cysteamine solutions. All recordings were voltage-clamped at −60 mV and data collected at 5 kHz using an Axopatch 200B amplifier and Digidata 1440A (Molecular Devices) with Axopatch software. Current decay was fit with a double exponential, and weighted time constants were obtained using:

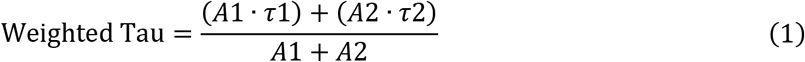

Where A1 and A2 are the amplitudes of the first and second exponential components. Peak responses as a function of cysteamine were fit to a Hill equation with n =2 to derive an EC_50_.

Statistical analysis was performed using multiple comparisons with a Dunnett’s test to compare fatty acid or KK-242 inhibition with control.

### Photolabeling of ELIC and intact protein MS

Photolabeling of ELIC by pacFA and KK-242 and intact protein MS analysis was performed as previously described (21). 50 μg of purified ELIC in buffer A was mixed with 100 μM pacFA or KK-242 for 30 min, and irradiated with >320 nm UV light for 5 min. For intact-protein MS analysis, 25 μg of photolabeled ELIC was precipitated with chloroform, methanol and water, pelleted and reconstituted in 3 μl of 100% formic acid for 10 s followed by 50 μl of 4:4:1 chloroform/methanol/water. These samples were analyzed in the ion trap (LTQ) of an Elite mass spectrometer (Thermo Scientific) by direct injection using a Max Ion API source with a HESI-II probe. Data was acquired with a flow rate of 3 μl/min, spray voltage of 4 kV, capillary temperature of 350 °C, and SID of 30 V. Spectra were deconvoluted using Unidec (50).

### Middle-down MS analysis of photolabeled ELIC

To sequence photolabeled ELIC, a tryptic middle-down MS analysis was performed as previously described with some modifications (21). 15 μg of KK-242 photolabeled ELIC was buffer exchanged to 50 mM triethylammonium bicarbonate (TEABC) pH 7.5 and 0.02% DDM using Biospin gel filtration spin columns (Bio-Rad). These samples were then reduced and alkylated with triscarboxyethylphosphine (TCEP) and N-ethylmaleimide (NEM) followed by digestion with 2 μg of trypsin for 7 days at 4 ^o^C. Digestion was terminated with formic acid at 1%. The final sample volume was 100 μl, and 20 μl was analyzed by LC-MS using an in-house PLRP-S (Agilent) column and Orbitrap Elite mass spectrometer (Thermo Scientific). MS/MS was performed as previously reported (37), using HCD (higher-energy collisional dissociation). The LC-MS data was searched with PEAKS (Bioinformatics Solutions) using the following search parameters: precursor mass accuracy of 20 ppm, fragment ion accuracy of 0.1 Da, up to three missed cleavages on either end of the peptide, false discovery rate of 0.1%, and variable modifications of methionine oxidation, cysteine alkylation with NEM, and the KK-242 mass (330.14 Da). Manual data analysis of MS1 and MS2 spectra was performed with XCalibur (Thermo Scientific). Photolabeled peptides were only accepted if there was a corresponding unlabeled peptide with a shorter a retention time, and a mass accuracy of <10 ppm. All fragment ions for MS2 spectra had a mass accuracy of <20 ppm.

To assess competition of KK-242 photolabeling by fatty acids, 15 μg of ELIC was photolabeled with 10 μM KK-242 in the absence and presence of different concentrations of DHA and PA. The photolabeling efficiency of M4 was obtained by taking the area under the curve of extracted ion chromatograms of unlabeled and labeled M4 in XCalibur. Photolabeling efficiency was calculated as the abundance (area under the curve) of labeled peptide/(unlabeled+labeled peptide).

### hMTS modification of ELIC mutants and reconstitution in giant liposomes

hMTS modification of ELIC was performed by mixing ELIC in buffer A at 1 mg/ml with equal volumes of 200 μM hMTS in buffer A for a final concentration of 100 μM hMTS and 0.5 mg/ml ELIC. This sample was incubated at RT for 30 min, followed by removal of 25 μg of protein for intact protein MS analysis and purification of the remaining protein on a Sephadex 200 Increase 10/300 column. For hMTS-modified R123C/C300S/C313S, this yielded a monodisperse protein, and 0.5 mg was reconstituted in giant liposomes in the same way as WT or C300S/C313S. Giant liposomes of hMTS-modified R117C/C300S/C313S were prepared as follows. 0.5 mg of purified protein was added to DDM-destabilized liposomes at 10 mg/ml (sample volume of 0.5 ml). After equilibrating for 30 min at RT, 0.5 ml of 200 μM hMTS in buffer A was added three times with each addition separated by 15 min (final volume 2 ml). Next, Bio-Beads were added to the remaining sample to remove DDM, as per the giant liposome formation protocol. SDS-PAGE analysis of ELIC protein from proteoliposomes was performed by solubilizing 2 μl of proteoliposomes (from frozen aliquots) in 1% SDS for 1 h at RT. Samples were then run on 10% Tris-glycine polyacrylamide gel (Bio-Rad), and protein was detected using SYPRO-Ruby staining (Thermo Fisher) and fluorescence imaging of the gel.

### Docking of KK-242 to photolabeled sites in ELIC

Docking was performed using Autodock 4.2 as previously described (21,51). The structure of KK-242 was generated in Maestro (Schrödinger), and Gasteiger charges and free torsion angles were determined by Autodock Tools. 2YN6 was used as the docking template, and two runs were performed with grids encompassing the two KK-242 photolabeled sites. The grid dimensions were 26 x 16 x 20 Å with 1 Å spacing for both sites and were centered to encompass the outer TMD interface and C313 or Q264 for each site. Each docking run produced 300 poses that were clustered at 2.0 Å RMSD. Poses selected for Figure 4 were taken from a cluster where the diazirine is closest to the photolabeled residue.

### Coarse-grained simulations of ELIC with fatty acid

For coarse-grained (CG) molecular dynamics simulations, the propylamine-bound non-conducting structure of ELIC (PDB 6V03) was used as the structural model (52). As the deposited PDB terminates at residue 317, residues 318-321 (318-RGIT-321) were modelled assuming M4 retained an α-helical nature until termination. The pKa of each sidechain was measured using PROPKA (53) and protonated assuming a pH of 7. The resulting ELIC structure was embedded and oriented in a POPC membrane using the CHARMM-GUI application (54,55). CHARMM-GUI was also utilized to solvate and ionize the system with 150 mM NaCl. The final system contained approximately 192,000 atoms and measured 122×122×143 Å^3^. To allow the additional residues on the M4 helix to relax, an equilibration of the all-atom system was performed. The simulation system was energy minimized for 10,000 steps. While keeping the protein backbone harmonically restrained (*k* = 5 kcal•mol^-1^•Å^-2^), the system was equilibrated for 5.0 ns in an NPT ensemble, allowing the membrane and aqueous phase to relax. The harmonic restraints on the protein backbone were then gradually released over a period of 5.0 ns under an NPT ensemble, after which the whole system was equilibrated without restraints for 5.0 ns. All atomistic simulations were performed with NAMD 2.14 (56) with CHARMM36 parameters (57,58). Temperature was maintained at 310K using a Langevin damping coefficient, y, of 1 ps^-1^. Pressure was maintained at 1.0 atm using the Nosé-Hoover Langevin piston method. Long-range electrostatic interactions were treated with particle mesh Ewald sums on a grid density > 1 Å^-3^. A timestep of 2.0 fs was used for all atomistic simulations with bonded and non-bonded forces calculated every timestep and particle mesh Ewald sums calculated every other timestep. Long-range non-bonded interactions were cutoff after 12.0 Å with a smoothing function applied between 10.0 Å and 12.0 Å.

The resulting equilibrated ELIC structure was coarse-grained using the *martinize* script (59), including secondary structural restraints. Conformational structure was maintained via harmonic restraints for backbone beads <0.5 nm apart using a force constant of 1000 kJ mol^-1^ with backbone pairs generated using the ElNeDyn algorithm (60). The coarse-grained ELIC structure was embedded in the membrane using the *insane* script (61). Each simulation system used a bulk membrane composed of 2:1:1 POPC:POPE:POPG to which either 4 mol% palmitic acid or 4 mol% docosahexaenoic acid was added. For each system simulated, this equated to 15 fatty acid molecules per leaflet, or 30 total for the system. After solvation and ionization to 150 mM NaCl, final system size was approximately 16×16×20 nm^3^ with approximately 44,000 beads.

Two initial simulation systems, one containing PA and one containing DHA, were created and equilibrated as below. Each system was energy minimized using steepest descent for approximately 30,000 steps and then equilibrated for 6.0 ns. The system was equilibrated in a NVT ensemble for 1.0 ns using the Berendsen thermostat set at 323K and a temperature coupling constant of 1.0 ps. Following this, the system was equilibrated in a NPT ensemble for 5.0 ns using the Berendsen thermostat and barostat. The temperature was set at 323K with a temperature coupling constant of 1.0 ps; the pressure was maintained at 1.0 bar using a pressure coupling constant of 3.0 ps and compressibility of 3.0×10^-5^ bar^-1^.

Once the initial systems were equilibrated, four replicates of each experimental condition were simulated for 10 μs, meaning a total of 40 μs of simulation data for PA and DHA each. For production simulations, temperature was maintained at 323K using the velocity rescaling algorithm (62) with the coupling constant set to 1.0 ps. The protein and lipid beads were coupled to a temperature bath separate from water and ions. Each bath had the same temperature parameters. Pressure was maintained at 1.0 bar semi-isotropically utilizing a time constant of 12 ps and a compressibility of 3.0×10^-5^ bar^-1^ with the Parinello-Rahman coupling scheme. All coarse-grained simulations were performed with GROMACS 2020.4 (63) and the MARTINI 2.2 force field (59,61). Electrostatics were treated with a reaction-field and dielectric constant of ε=15; van der Waals and Coulomb interactions were cut-off after 1.1 nm. A timestep of 25 fs was used throughout. Frames were saved every 1.0 ns and used for analysis as described below.

To understand the boundary distribution of fatty acids, the boundary lipid metric, B, was measured according to:

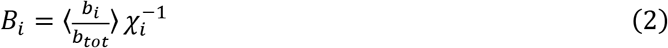

where *b_i_* is the number of lipids of species *i* in the boundary shell, *b_tot_* is the total number of lipids in the boundary shell, and *χ*_i_ is the fraction of lipid species *i* in the model membrane. The brackets denote an average over the time of the simulation. A lipid was considered in the boundary shell if it was within 6 Å of the protein.

Two-dimensional radial enhancement calculations were performed as demonstrated previously (30,35) to characterize fatty acid interactions with ELIC. The two-dimensional radial density distribution, ρ_B_, was calculated according to:

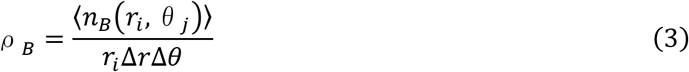

where ‹*n_B_*(*r_i_, θ_j_*)>› is the time-averaged number of beads of lipid species *B* in the bin centered around radius *r_i_* and polar angle *θ_j_, Δr* is the radial bin width (5 Å in this study), and *Δθ* is the polar angle bin width (π/15 radians in this study). In order to determine relative enrichment or depletion of the lipid compared to the bulk membrane phase, the two-dimensional radial density distribution was normalized to expected bead density of lipid species *B* in the bulk:

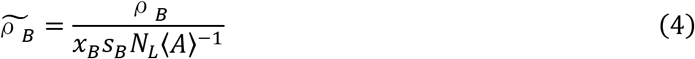

where *x_B_* is the mol fraction of lipid species *B, s_B_* is the number of coarse-grained beads in lipid species *B, N_L_* is the total number of lipids in the simulation system, and ‹*A*› is the average projected area of the simulation box in the plane of the membrane.

Contact probabilities of each individual residue with fatty acids were determined as follows. In each simulation frame, if any fatty acid bead was located within 5 Å of any bead of the residue, a contact was counted for that frame. The number of contacts was then time-averaged over the simulation and then averaged across all five subunits.

A density affinity free energy analysis was performed as previously described *(35)*. Briefly, this analysis determines the relative free energy difference for a lipid species between the bulk membrane and a particular binding site by comparing the probability distribution of finding *n* coarse-grained fatty acid beads in the binding site, *P_site_(n)*, versus the same probability distribution for bulk membrane, *P_bulk_(n)*. In this study, we defined two new binding sites, termed “M1” and “M3” (Fig. 2 and Fig. 5C). For the M1 site, one angular boundary was determined to be the M4 helix and the other angular boundary to be midway between the M1 helix of the same subunit and the M3 helix of the adjacent subunit. For the M3 site, one angular boundary was determined to be the M4 helix and the other angular boundary to be midway between the M3 helix of the same subunit and the M1 helix of the adjacent subunit. The radial boundaries for both sites were 10 < *r* < 40 Å. *P_site_(n)* was determined by counting the number of coarse-grained fatty acid beads within the boundaries of each binding site described above (*i.e*., M1 and M3) for each frame and normalizing to the number of frames in the simulation. As *P_bulk_(n)* needs to be corrected for the area of *P_site_(n)*, a separate simulation was conducted to determine the area of the M1 and M3 sites (35). Exploiting the relationship that *P_site_(n)* = *P_bulk_(n)* in a unitary membrane, a coarse-grained system containing ELIC (PDB 6V03) in a POPC membrane was simulated under the same conditions described above for 10 μs.

### Macroscopic current simulations with Channelab

Simulations of current responses were performed using Channelab utilizing a pLGIC gating model as previously described (34). Currents were simulated using the model and rate constants indicated in Figure 5 Supplement 3 and 4, and a fourth-order Runge-Kutta integration algorithm. Each simulation was run pre-equilibrated in the presence of varying concentrations of inhibitor, followed by the application of agonist for 25 s. Channel currents were determined by monitoring the probability of occupying the AO or AOI states. Occupancy of pre-active or desensitized states were monitored by calculating the probability of occupying the AF/AFI or AD/ADI states, respectively.

## ACKNOWLEDGEMENTS

This work was funded by grants from the National Institutes of Health (NIH) including R35GM137957 for W.W.C., F32GM139351 for J.T.P., and R01HL067773 and R01GM108799 for D.F.C. We are grateful to Alex S. Evers, Gustav Akk, and Joe Henry Steinbach for helpful discussions and comments on the manuscript. We are grateful to Stephen Traynelis for sharing the Channelab software.

## CONFLICTS OF INTEREST

The authors declare that they have no conflicts of interest with the contents of the article.

## AUTHOR CONTRIBUTIONS

N.D., M.J.A., L.J.C. and J.T.P. contributed to the acquisition and analysis of data and writing of the manuscript. K.K. contributed to the synthesis of KK-242. G.B. contributed to the study design, analysis of data and writing of the manuscript. D.F.C. contributed to the study design and synthesis of KK-242. W.W.C. contributed to the study conception and design, analysis of data, and writing of the manuscript.

## Notes

### Competing Interest Statement

The authors have declared no competing interest.

## REFERENCES

1. Antollini, S. S., and Barrantes, F. J. (2016) Fatty Acid Regulation of Voltage- and Ligand-Gated Ion Channel Function. Front Physiol 7, 573

2. Fernandez Nievas, G. A., Barrantes, F. J., and Antollini, S. S. (2008) Modulation of nicotinic acetylcholine receptor conformational state by free fatty acids and steroids. J Biol Chem 283, 21478–21486

3. Basak, S., Schmandt, N., Gicheru, Y., and Chakrapani, S. (2017) Crystal structure and dynamics of a lipid-induced potential desensitized-state of a pentameric ligand-gated channel. Elife 6

4. Nabekura, J., Noguchi, K., Witt, M. R., Nielsen, M., and Akaike, N. (1998) Functional modulation of human recombinant gamma-aminobutyric acid type A receptor by docosahexaenoic acid. J Biol Chem 273, 11056–11061

5. Hamano, H., Nabekura, J., Nishikawa, M., and Ogawa, T. (1996) Docosahexaenoic acid reduces GABA response in substantia nigra neuron of rat. J Neurophysiol 75, 1264–1270

6. Minota, S., and Watanabe, S. (1997) Inhibitory effects of arachidonic acid on nicotinic transmission in bullfrog sympathetic neurons. J Neurophysiol 78, 2396–2401

7. McNamara, R. K., and Carlson, S. E. (2006) Role of omega-3 fatty acids in brain development and function: potential implications for the pathogenesis and prevention of psychopathology. Prostaglandins Leukot Essent Fatty Acids 75, 329–349

8. Hashimoto, M., Hossain, S., Al Mamun, A., Matsuzaki, K., and Arai, H. (2017) Docosahexaenoic acid: one molecule diverse functions. Crit Rev Biotechnol 37, 579–597

9. Taha, A. Y., Filo, E., Ma, D. W., and McIntyre Burnham, W. (2009) Dose-dependent anticonvulsant effects of linoleic and alpha-linolenic polyunsaturated fatty acids on pentylenetetrazol induced seizures in rats. Epilepsia 50, 72–82

10. Taha, A. Y., Zahid, T., Epps, T., Trepanier, M. O., Burnham, W. M., Bazinet, R. P., and Zhang, L. (2013) Selective reduction of excitatory hippocampal sharp waves by docosahexaenoic acid and its methyl ester analog ex-vivo. Brain Res 1537, 9–17

11. Cordero-Morales, J. F., and Vasquez, V. (2018) How lipids contribute to ion channel function, a fat perspective on direct and indirect interactions. Curr Opin Struct Biol 51, 92–98

12. Sogaard, R., Werge, T. M., Bertelsen, C., Lundbye, C., Madsen, K. L., Nielsen, C. H., and Lundbaek, J. A. (2006) GABA(A) receptor function is regulated by lipid bilayer elasticity. Biochemistry 45, 13118–13129

13. Tian, Y., Aursnes, M., Hansen, T. V., Tungen, J. E., Galpin, J. D., Leisle, L., Ahern, C. A., Xu, R., Heinemann, S. H., and Hoshi, T. (2016) Atomic determinants of BK channel activation by polyunsaturated fatty acids. Proc Natl Acad Sci U S A 113, 13905–13910

14. Borjesson, S. I., Hammarstrom, S., and Elinder, F. (2008) Lipoelectric modification of ion channel voltage gating by polyunsaturated fatty acids. Biophys J 95, 2242–2253

15. Yazdi, S., Stein, M., Elinder, F., Andersson, M., and Lindahl, E. (2016) The Molecular Basis of Polyunsaturated Fatty Acid Interactions with the Shaker Voltage-Gated Potassium Channel. PLoS Comput Biol 12, e1004704

16. Haberkant, P., and Holthuis, J. C. (2014) Fat & fabulous: bifunctional lipids in the spotlight. Biochim Biophys Acta 1841, 1022–1030

17. Haberkant, P., Raijmakers, R., Wildwater, M., Sachsenheimer, T., Brugger, B., Maeda, K., Houweling, M., Gavin, A. C., Schultz, C., van Meer, G., Heck, A. J., and Holthuis, J. C. (2013) In vivo profiling and visualization of cellular protein-lipid interactions using bifunctional fatty acids. Angew Chem Int Ed Engl 52, 4033–4038

18. Capone, J., Leblanc, P., Gerber, G. E., and Ghosh, H. P. (1983) Localization of membrane proteins by the use of a photoreactive fatty acid incorporated in vivo into vesicular stomatitis virus. J Biol Chem 258, 1395–1398

19. Budelier, M. M., Cheng, W. W. L., Chen, Z. W., Bracamontes, J. R., Sugasawa, Y., Krishnan, K., Mydock-McGrane, L., Covey, D. F., and Evers, A. S. (2019) Common binding sites for cholesterol and neurosteroids on a pentameric ligand-gated ion channel. Biochim Biophys Acta Mol Cell Biol Lipids 1864, 128–136

20. Budelier, M. M., Cheng, W. W. L., Bergdoll, L., Chen, Z.W., Abramson, J., Krishnan, K., Qian, M, Covey, D.F., Janetka, J.W., and Evers, A.S. (2017) Click chemistry reagent for identification of sites of covalent ligand incorporation in integral membrane proteins. Analytical Chemistry

21. Cheng, W. W. L., Chen, Z. W., Bracamontes, J. R., Budelier, M. M., Krishnan, K., Shin, D. J., Wang, C., Jiang, X., Covey, D. F., Akk, G., and Evers, A. S. (2018) Mapping two neurosteroid-modulatory sites in the prototypic pentameric ligand-gated ion channel GLIC. J Biol Chem 293, 3013–3027

22. Hilf, R. J., and Dutzler, R. (2008) X-ray structure of a prokaryotic pentameric ligand-gated ion channel. Nature 452, 375–379

23. Henault, C. M., Govaerts, C., Spurny, R., Brams, M., Estrada-Mondragon, A., Lynch, J., Bertrand, D., Pardon, E., Evans, G. L., Woods, K., Elberson, B. W., Cuello, L. G., Brannigan, G., Nury, H., Steyaert, J., Baenziger, J. E., and Ulens, C. (2019) A lipid site shapes the agonist response of a pentameric ligand-gated ion channel. Nat Chem Biol 15, 1156–1164

24. Tong, A., Petroff, J. T., 2nd, Hsu, F. F., Schmidpeter, P. A., Nimigean, C. M., Sharp, L., Brannigan, G., and Cheng, W. W. (2019) Direct binding of phosphatidylglycerol at specific sites modulates desensitization of a ligand-gated ion channel. Elife 8

25. Kinde, M. N., Bu, W., Chen, Q., Xu, Y., Eckenhoff, R. G., and Tang, P. (2016) Common Anesthetic-binding Site for Inhibition of Pentameric Ligand-gated Ion Channels. Anesthesiology 124, 664–673

26. Chen, Q., Kinde, M. N., Arjunan, P., Wells, M. M., Cohen, A. E., Xu, Y., and Tang, P. (2015) Direct Pore Binding as a Mechanism for Isoflurane Inhibition of the Pentameric Ligand-gated Ion Channel ELIC. Sci Rep 5, 13833

27. Spurny, R., Billen, B., Howard, R. J., Brams, M., Debaveye, S., Price, K. L., Weston, D.A., Strelkov, S. V., Tytgat, J., Bertrand, S., Bertrand, D., Lummis, S. C., and Ulens, C. (2013) Multisite binding of a general anesthetic to the prokaryotic pentameric Erwinia chrysanthemi ligand-gated ion channel (ELIC). J Biol Chem 288, 8355–8364

28. Laha, K. T., Ghosh, B., and Czajkowski, C. (2013) Macroscopic kinetics of pentameric ligand gated ion channels: comparisons between two prokaryotic channels and one eukaryotic channel. PLoS One 8, e80322

29. Vijayaraghavan, S., Huang, B., Blumenthal, E. M., and Berg, D. K. (1995) Arachidonic acid as a possible negative feedback inhibitor of nicotinic acetylcholine receptors on neurons. J Neurosci 15, 3679–3687

30. Sharp, L., Salari, R., and Brannigan, G. (2019) Boundary lipids of the nicotinic acetylcholine receptor: Spontaneous partitioning via coarse-grained molecular dynamics simulation. Biochim Biophys Acta Biomembr 1861, 887–896

31. Sugasawa, Y., Bracamontes, J. R., Krishnan, K., Covey, D. F., Reichert, D. E., Akk, G., Chen, Q., Tang, P., Evers, A. S., and Cheng, W. W. L. (2019) The molecular determinants of neurosteroid binding in the GABA(A) receptor. J Steroid Biochem Mol Biol 192, 105383

32. Budelier, M. M., Cheng, W. W. L., Bergdoll, L., Chen, Z. W., Janetka, J. W., Abramson, J., Krishnan, K., Mydock-McGrane, L., Covey, D. F., Whitelegge, J. P., and Evers, A. S. (2017) Photoaffinity labeling with cholesterol analogues precisely maps a cholesterol-binding site in voltage-dependent anion channel-1. J Biol Chem 292, 9294–9304

33. Kumar, P., Cymes, G. D., and Grosman, C. (2021) Structure and function at the lipid-protein interface of a pentameric ligand-gated ion channel. Proc Natl Acad Sci U S A 118

34. Gielen, M., and Corringer, P. J. (2018) The dual-gate model for pentameric ligand-gated ion channels activation and desensitization. J Physiol 596, 1873–1902

35. Sharp, L., and Brannigan, G. (2021) Spontaneous lipid binding to the nicotinic acetylcholine receptor in a native membrane. J Chem Phys 154, 185102

36. Kinde, M. N., Chen, Q., Lawless, M. J., Mowrey, D. D., Xu, J., Saxena, S., Xu, Y., and Tang, P. (2015) Conformational Changes Underlying Desensitization of the Pentameric Ligand-Gated Ion Channel ELIC. Structure 23, 995–1004

37. Sugasawa, Y., Cheng, W. W., Bracamontes, J. R., Chen, Z. W., Wang, L., Germann, A. L., Pierce, S. R., Senneff, T. C., Krishnan, K., Reichert, D. E., Covey, D. F., Akk, G., and Evers, A. S. (2020) Site-specific effects of neurosteroids on GABAA receptor activation and desensitization. Elife 9

38. Lee, S. J., Wang, S., Borschel, W., Heyman, S., Gyore, J., and Nichols, C. G. (2013) Secondary anionic phospholipid binding site and gating mechanism in Kir2.1 inward rectifier channels. Nat Commun 4, 2786

39. Enkvetchakul, D., Jeliazkova, I., Bhattacharyya, J., and Nichols, C. G. (2007) Control of inward rectifier K channel activity by lipid tethering of cytoplasmic domains. J Gen Physiol 130, 329–334

40. Sridhar, A., Lummis, S. C. R., Pasini, D., Mehregan, A., Brams, M., Kambara, K., Bertrand, D., Lindahl, E., Howard, R. J., and Ulens, C. (2021) Regulation of a pentameric ligandgated ion channel by a semiconserved cationic lipid-binding site. J Biol Chem 297, 100899

41. Farag, N. E., Jeong, D., Claydon, T., Warwicker, J., and Boyett, M. R. (2016) Polyunsaturated fatty acids inhibit Kv1.4 by interacting with positively charged extracellular pore residues. Am J Physiol Cell Physiol 311, C255–268

42. Yazdi, S., Nikesjo, J., Miranda, W., Corradi, V., Tieleman, D. P., Noskov, S. Y., Larsson, H. P., and Liin, S. I. (2021) Identification of PUFA interaction sites on the cardiac potassium channel KCNQ1. J Gen Physiol 153

43. Cheng, W. W., D’Avanzo, N., Doyle, D. A., and Nichols, C. G. (2011) Dual-mode phospholipid regulation of human inward rectifying potassium channels. Biophys J 100, 620–628

44. Hite, R. K., Butterwick, J. A., and MacKinnon, R. (2014) Phosphatidic acid modulation of Kv channel voltage sensor function. Elife 3

45. Gao, Y., Cao, E., Julius, D., and Cheng, Y. (2016) TRPV1 structures in nanodiscs reveal mechanisms of ligand and lipid action. Nature 534, 347–351

46. Hansen, S. B., Tao, X., and MacKinnon, R. (2011) Structural basis of PIP2 activation of the classical inward rectifier K+ channel Kir2.2. Nature 477, 495–498

47. Simard, J. R., Zunszain, P. A., Ha, C. E., Yang, J. S., Bhagavan, N. V., Petitpas, I., Curry, S., and Hamilton, J. A. (2005) Locating high-affinity fatty acid-binding sites on albumin by x-ray crystallography and NMR spectroscopy. Proc Natl Acad Sci U S A 102, 17958–17963

48. Dadsena, S., Bockelmann, S., Mina, J. G. M., Hassan, D. G., Korneev, S., Razzera, G., Jahn, H., Niekamp, P., Muller, D., Schneider, M., Tafesse, F. G., Marrink, S. J., Melo, M. N., and Holthuis, J. C. M. (2019) Ceramides bind VDAC2 to trigger mitochondrial apoptosis. Nat Commun 10, 1832

49. Hulce, J. J., Cognetta, A. B., Niphakis, M. J., Tully, S. E., and Cravatt, B. F. (2013) Proteome-wide mapping of cholesterol-interacting proteins in mammalian cells. Nat Methods 10, 259–264

50. Marty, M. T., Baldwin, A. J., Marklund, E. G., Hochberg, G. K., Benesch, J. L., and Robinson, C. V. (2015) Bayesian deconvolution of mass and ion mobility spectra: from binary interactions to polydisperse ensembles. Anal Chem 87, 4370–4376

51. Morris, G. M., Huey, R., Lindstrom, W., Sanner, M. F., Belew, R. K., Goodsell, D. S., and Olson, A. J. (2009) AutoDock4 and AutoDockTools4: Automated docking with selective receptor flexibility. J Comput Chem 30, 2785–2791

52. Kumar, P., Wang, Y., Zhang, Z., Zhao, Z., Cymes, G. D., Tajkhorshid, E., and Grosman, C. (2020) Cryo-EM structures of a lipid-sensitive pentameric ligand-gated ion channel embedded in a phosphatidylcholine-only bilayer. Proc Natl Acad Sci U S A 117, 1788–1798

53. Olsson, M. H., Sondergaard, C. R., Rostkowski, M., and Jensen, J. H. (2011) PROPKA3: Consistent Treatment of Internal and Surface Residues in Empirical pKa Predictions. J Chem Theory Comput 7, 525–537

54. Jo, S., Kim, T., Iyer, V. G., and Im, W. (2008) CHARMM-GUI: a web-based graphical user interface for CHARMM. J Comput Chem 29, 1859–1865

55. Wu, E. L., Cheng, X., Jo, S., Rui, H., Song, K. C., Davila-Contreras, E. M., Qi, Y., Lee, J., Monje-Galvan, V., Venable, R. M., Klauda, J. B., and Im, W. (2014) CHARMM-GUI Membrane Builder toward realistic biological membrane simulations. J Comput Chem 35, 1997–2004

56. Phillips, J. C., Hardy, D. J., Maia, J. D. C., Stone, J. E., Ribeiro, J. V., Bernardi, R. C., Buch, R., Fiorin, G., Henin, J., Jiang, W., McGreevy, R., Melo, M. C. R., Radak, B. K., Skeel, R. D., Singharoy, A., Wang, Y., Roux, B., Aksimentiev, A., Luthey-Schulten, Z., Kale, L. V., Schulten, K., Chipot, C., and Tajkhorshid, E. (2020) Scalable molecular dynamics on CPU and GPU architectures with NAMD. J Chem Phys 153, 044130

57. Klauda, J. B., Venable, R. M., Freites, J. A., O’Connor, J. W., Tobias, D. J., Mondragon-Ramirez, C., Vorobyov, I., MacKerell, A. D., Jr., and Pastor, R. W. (2010) Update of the CHARMM all-atom additive force field for lipids: validation on six lipid types. J Phys Chem B 114, 7830–7843

58. Huang, J., Rauscher, S., Nawrocki, G., Ran, T., Feig, M., de Groot, B. L., Grubmuller, H., and MacKerell, A. D., Jr. (2017) CHARMM36m: an improved force field for folded and intrinsically disordered proteins. Nat Methods 14, 71–73

59. de Jong, D. H., Singh, G., Bennett, W. F., Arnarez, C., Wassenaar, T. A., Schafer, L. V., Periole, X., Tieleman, D. P., and Marrink, S. J. (2013) Improved Parameters for the Martini Coarse-Grained Protein Force Field. J Chem Theory Comput 9, 687–697

60. Periole, X., Cavalli, M., Marrink, S. J., and Ceruso, M. A. (2009) Combining an Elastic Network With a Coarse-Grained Molecular Force Field: Structure, Dynamics, and Intermolecular Recognition. J Chem Theory Comput 5, 2531–2543

61. Wassenaar, T. A., Ingolfsson, H. I., Bockmann, R. A., Tieleman, D. P., and Marrink, S. J. (2015) Computational Lipidomics with insane: A Versatile Tool for Generating Custom Membranes for Molecular Simulations. J Chem Theory Comput 11, 2144–2155

62. Bussi, G., Donadio, D., and Parrinello, M. (2007) Canonical sampling through velocity rescaling. J Chem Phys 126, 014101

63. Kutzner, C., Pall, S., Fechner, M., Esztermann, A., de Groot, B. L., and Grubmuller, H. (2019) More bang for your buck: Improved use of GPU nodes for GROMACS 2018. J Comput Chem 40, 2418–2431

